# Perturbation of mitochondrial Ca^2+^ homeostasis activates cross-compartmental proteostatic response in Arabidopsis

**DOI:** 10.1101/2022.10.02.510489

**Authors:** Xiaoyan Zhang, Chongyang Ma, Xinyue Bao, Shenyu Zhang, Omar Zayed, Zhengjing Zhang, Kai Tang, Shaojun Xie, Yunsheng Wang, Dayong Zhang, Huawei Xu, Huifang Jia, Xinying Wang, Qianyan Lei, Xiaocui Wang, Junli Zhang, Savithramma P Dinesh-Kumar, Chun-Peng Song, Jian-Kang Zhu, Xiaohong Zhu

## Abstract

Mitochondrial Ca^2+^ (_mt_Ca^2+^) homeostasis is essential to mitochondrial functions. However, how _mt_Ca^2+^ homeostasis is achieved and the consequences of impaired _mt_Ca^2+^ homeostasis in plants is poorly understood. Here, we demonstrate a critical role for mitochondrial Ca^2+^ uniporter (MCU) in the control of _mt_Ca^2+^ uptake for _mt_Ca^2+^ homeostasis *in planta* by characterizing MCU mutants and overexpressed plants. Impaired MCU-controlled _mt_Ca^2+^ homeostasis (iMUCH) in gain-of-function and loss-of-function MCU plants causes the misregulation of mitochondrial gene expression that triggers mitonuclear protein imbalance. Transcriptome integrated with proteomics analysis reveal activation of multiple compartmental UPR gene expression and decrease of cytosolic translation with selective repression of ribosome and RNA modification protein synthesis upon iMUCH. Intriguingly, TOR signalling is not involved in cytosolic translational response to iMUCH, but the reduction of eIFα phosphorylation is evident under iMUCH induced mitochondrial stress. Thus, our study unveils the essential functions of MCU proteins for _mt_Ca^2+^ homeostasis, and the involvement of MCU-controlled _mt_Ca^2+^ homeostasis in mitochondrial stress dependent regulation of protein synthesis for cellular proteostasis that is connected to plant growth and stress resistance.

## Introduction

Mitochondria are multifunctional organelles that host key biochemical pathways for energy conversion, as well as amino acid and lipid metabolism. Mitochondria also play a critical signaling role in aging (senescence), programmed cell death (PCD), and stress responses. They directly perceive external stress stimuli, including extreme temperatures, drought, and high salinity, which adversely influence mitochondrial function and trigger mitochondrial stress ^1,2^. To cope with these stress challenges, mitochondria are in constant communication with the nucleus and the cytosol to reprogram nuclear transcription, and to reset the cytosolic translation machinery, which are well characterized by the mitochondria-to-nucleus communication ^3-7^ and the mitochondrial-to-cytosolic response ^8-12^. In animal cells, the mitochondrial unfolded protein response (UPR^mt^) is induced by an imbalance in protein translation between nucleus-encoded and mitochondrion-encoded proteins. UPR^mt^ is a key mitochondrial retrograde signaling pathway that ensures proper translation, folding, and degradation of mitochondrion-localized proteins ^13-15^. Whether these mitochondrial stress adaptive responses are conserved in plants and, if so, whether they share common players and biological consequences with their animal counterparts remain unclear.

The mitochondrial matrix Ca^2+^ (_mt_Ca^2+^) fluctuates in response to environmental and/or cellular cues ^16,17^. Disruption of _mt_Ca^2+^ homeostasis sensitizes mitochondria to the challenges of stress, aging (senescence), and apoptosis ^18-23^. However, it is not fully understood how disturbed _mit_Ca^2+^ homeostasis impairs mitochondrial function though it is acknowledged that mitochondrial Ca^2+^ uptake buffers cytosolic Ca^2+^ signaling ^17,24^. The mitochondrial Ca^2+^ uniporter protein complex (MCUC) in mammalian systems facilitates Ca^2+^ uptake through pore-forming MCUs whose activity is modulated by the regulatory components MITOCHONDRIAL Ca^2+^ UPTAKE (MICU) and ESSENTIAL MCU REGULATOR (EMRE) ^25-31^. Structural analysis of MCUC by cryo-electron microscopy revealed the gating mechanism by which MICUs control uniporter activity ^32^. The plant MICU also negatively fine-tunes MCU-mediated mitochondrial Ca^2+^ uptake, thus preserving _mt_Ca^2+^ homeostasis ^33^. Six putative *Arabidopsis thaliana* (Arabidopsis) MCU orthologs have been identified, each with a relatively conserved transmembrane domain, a pore loop, and the signature sequence DVME ^34^. Arabidopsis MCU1 (At1g09575), MCU2 (At1g57610), MCU3 (At2g23790), and MCU5 (At5g42610) appear to be localized to mitochondria, while MCU6 (At5g66650, CMCU) is targeted to both mitochondria and chloroplasts ^34-37^. Stress stimulus-specific Ca^2+^ dynamics in the chloroplast stroma (_cs_Ca^2+^) are correlated with the transcript levels of *MCU6* ^36^. A recent study reported that mitochondrial Ca^2+^ uptake became limiting in the root cell of *mcu1 mcu2 mcu3* triple mutant, demonstrating that MCU proteins mediate mitochondrial Ca^2+^ uptake *in vivo* ^38^.

To study whether MCU proteins are necessary for maintaining _mt_Ca^2+^ homeostasis *in planta*, and how impaired MCU-controlled _mt_Ca^2+^ homeostasis affects mitochondrial function, we generated single and sextuple loss-of-function mutants of *MCU*s, as well as stable transgenic plants overexpressing *MCUs*, respectively. We show that mitochondrial Ca^2+^ uptake in response to stress stimuli are altered upon *MCU* overexpression and in sextuple mutant plants, demonstrating that MCUs are essential for _mt_Ca^2+^ homeostasis. We discover that impaired MCU-controlled _mt_Ca^2+^ homeostasis causes mitonuclear protein imbalance and activates proteostatic responses in multiple cellular compartments.

## Results

### Mitochondrial Ca^2+^ uniporters are necessary for _mt_Ca^2+^ homeostasis in plants

We first examined the subcellular localization of six putative MCUs ^34^(Teardo *et al*., 2017) in stable transgenic Arabidopsis plants expressing *MCU1, MCU2, MCU3, MCU4, MCU5*, or *MCU6* fused to yellow fluorescent protein (YFP) under the control of 35S promoter (*35S:MCU1-YFP* to *35S:MCU6-YFP*). YFP signal was predominantly observed in mobile mitochondrion-like structures in epidermal cells and mesophyll cells of the leaf (Supplementary Fig. 1A), and co-localized with the MitoTracker in root cells of these transgenic plants (Supplementary Fig. 1B), supporting MCUs localization to the mitochondria. Expression patterns of the 6 *MCU* genes cross tissues is important information to dissect their functions. We created individual transgenic Arabidopsis GUS reporter line driven by the *MCU* gene promoter (see Methods). GUS staining showed that the 6 *MCU* genes were all expressed in the stele of the root, and *MCU3* was highly expressed in the root apex including the root cap and quiescent centre, and in the root cortex. In contrast, *MCU6* was highly expressed in the basal meristem (Supplementary Fig. S1C, upper panel). *MCU3, MCU4*, and *MCU6*, but not *MCU1, MCU2* and *MCU5* were expressed in guard cells (Supplementary Fig. 1C, middle panel). All six *MCU* genes were expressed in anthers, however, *MCU1* and *MCU5* expression was relatively low compared to the other 4 MCUs (Supplementary Fig. 1C, lower panel). These results indicate that the six putative Arabidopsis MCUs are mitochondrial proteins with tissue-specific expression patterns.

Stress stimuli evoke cytosolic Ca^2+^ transient and mitochondrial Ca^2+^ uptake in plants ^16,17,36-42^. To study the function of MCUs in the control of _mt_Ca^2+^ homeostasis, we generated transgenic Arabidopsis Col-0 lines in which Ca^2+^ reporter *Aequorin* was targeted to mitochondrial matrix under the control of ubiquitin promoter (Col:*mtAq*). Next, we introduced CRISPR/Cas9 construct with two guide RNAs targeting each of the *MCU* genes (Supplementary Fig. 2A) in Col:*mtAq* line. We obtained the single copy of homozygous CAS9-edited *mcu* single mutants in Col:*mtAq* line, *mcu:mtAq* (Supplementary Fig. 2B). A significant reduction of *MCU* transcripts were detected in two generations of *mcu:mtAq* lines, respectively, indicating that CAS9-edited *MCUs* in the Col:*mtAq* reporter lines are effective and stable (Supplementary Fig. 2C). We used these *mcu:mtAeq* mutant lines and Col:*mtAq* to determine if MCU proteins are involved in mitochondrial Ca^2+^ uptake using luminometer. We observed similar _mt_Ca^2+^ dynamics with no differences for _mt_Ca^2+^ amplitudes in *mcu:mtAeq* mutants compared to Col:*mtAq* in response to mannitol, except for *mcu5:mtAq* showing an average reduction of 12% in _mt_Ca^2+^ amplitudes (Supplementary Fig. 3A and 3C). We detected no differences for _mt_Ca^2+^ amplitudes evoked by NaCl in single mutants except for an average reduction of 17% in *mcu3:mtAq*. These results indicate that loss of each member of MCU family have no major effects on _mt_Ca^2+^ amplitudes in response to tested stimuli.

We reasoned that lack of effect on _mt_Ca^2+^ amplitudes could be due to redundancy in the function of *MCU* genes. To overcome this, we expressed *mtAq* reporter construct into a *mcu* sextuple mutant that was generated by crossing *mcu* single mutants, *mcu1-1* (SALK_082151), *mcu2-1* (SALK_011710), *mcu3-1* (SALK_019312), *mcu4-1* (SALK_036975), *mcu6-1* (SALK_037347) and a CRISPR/Cas9-edited *mcu5-1* (Supplementary Fig. 2D and 2E) to obtain *mcu*^*1-6*^*:mtAq*. During our analysis, we discovered a deletion in the last exon of *MCU3* in the *mcu3-1* mutant, which is presumably caused by T-DNA insertion. This deletion likely results in a MCU3 protein with a C-terminal truncation (Supplementary Fig. 2G and 2H). *MCU* transcripts are significantly reduced in *mcu* sextuple mutant, indicating that it is a loss-of-function *mcu* sextuple mutants (Supplementary Fig. 2F and 2I). To examine how overexpression of MCU could affect _mt_Ca^2+^ homeostasis, we chose one of *MCU* overexpression lines, *2OX-1* that showed the high *MCU2* expression level compared to other lines (Supplementary Fig. 4, Fig. 7) to ensure an effectiveness of gain-of-function MCU. 2OX-1 was crossed to Col:*mtAq* to generate *2OX:mtAq* line. Transcript analysis revealed 9 fold increased expression of *MCU2* in *2OX:mtAq* (Supplementary Fig. 2I).

We used *2OX:mtAq* and *mcu*^*1-6*^*:mtAq* to determine the effect of overexpression and reduced expression of *MCU* on mitochondrial Ca^2+^ uptake in response to stimuli using Film Adhesive Seedling (FAS) imaging ^40^(Zhu *et al*., 2013) and luminometer reading. FAS imaging demonstrates that the _mt_Ca^2+^ in roots and leaves of *2OX:mtAq* seedlings showed strong response to mannitol and NaCl, but relatively weak response in *mcu*^*1-6*^*:mtAq* compared to that of Col:*mtAq*, respectively (Fig.1a and 1b). Although similar _mt_Ca^2+^ dynamics of *2OX:mtAq* and *mcu*^*1-6*^-*mtAq* were recorded by luminometer (Fig. 1c, d), the higher _mt_Ca^2+^ amplitudes and curve area were detected in *2OX:mtAq* compared to Col:*mtAq* plants in response to mannitol and NaCl (Fig. 1e-h), indicating that the mitochondrial Ca^2+^ uptake is increased by overexpression of MCU2. By contrast, the *mcu*^*1-6*^*:mtAq* displayed a significant reduction in mannitol- and NaCl-triggered _mt_Ca^2+^ amplitudes, an average reduction of 21% and 26% for mannitol and NaCl, respectively (Fig. 1e, g), and a slight reduction in curve area compared to Col:*mtAq* plants (Fig. 1f, h). Both *2OX:mtAq* and *mcu*^*1-6*^*:mtAq* plants showed a delay in peak time of _mt_Ca^2+^ response to NaCl (Fig. 1i). To determine if cytosolic Ca^2+^ is affected, we introduced cytosolic Aquorin reporter construct into wildtype (Col:*cytAq), 2OX-1* (*2OX:cytAq*) and *mcu*^*1-6*^ lines (*mcu*^*1-6*^*:cytAq*). We observed no significant differences in cytosolic Ca^2+^ between *2OX:cytAq* and Col:*cytAq* seedlings or between *mcu*^*1-6*^*:cytAq* and Col:*cytAq* seedlings in response to mannitol and NaCl (Fig. 1j, k). These results indicate that Arabidopsis MCUs play a role in the control of mitochondrial Ca^2+^ uptake for maintaining _mt_Ca^2+^ homeostasis with functional redundancy.

**Fig. 1.**
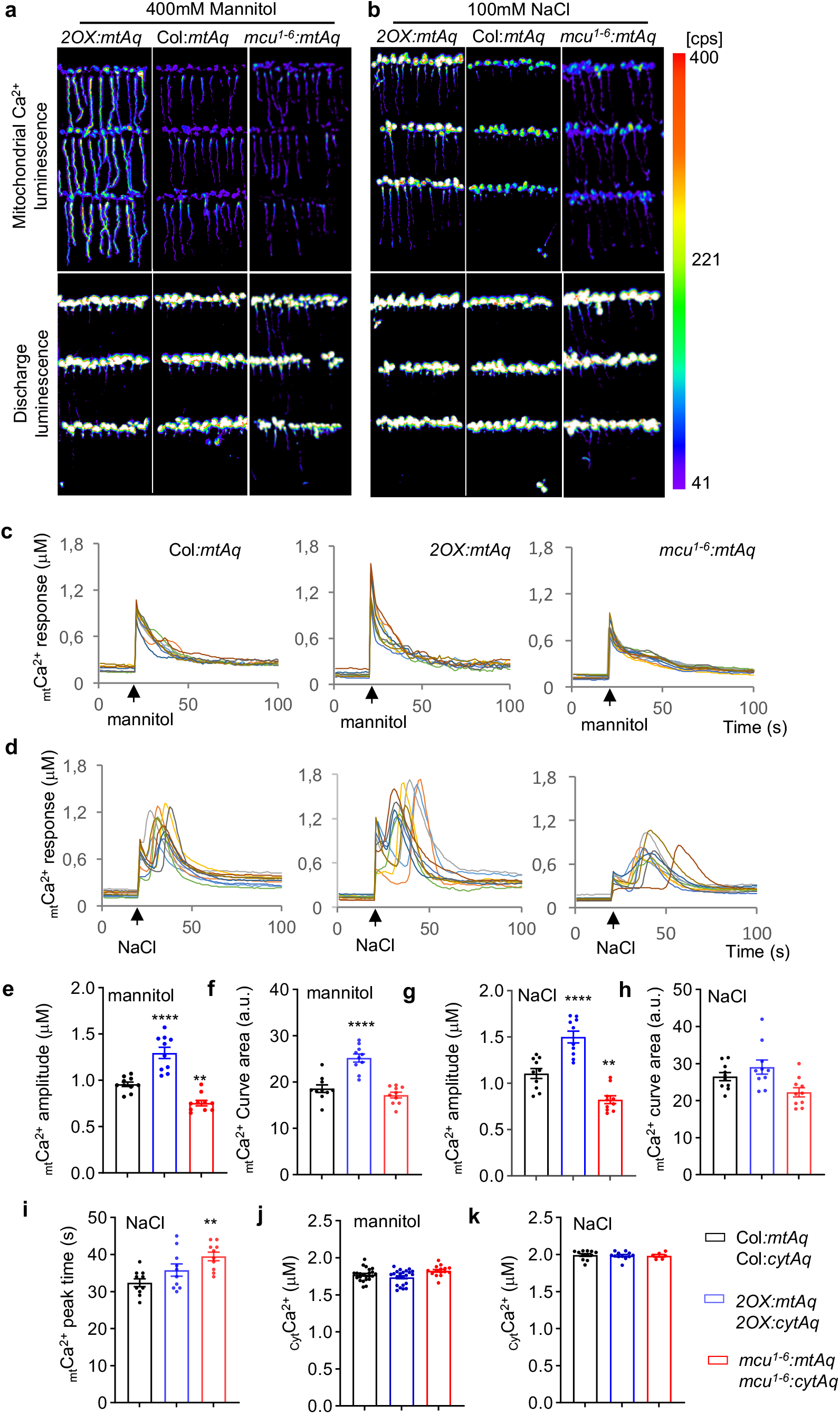
Altered *MCU*s expression affects mitochondrial Ca^2+^ uptake. **a, b** FAS imaging of Aequorin-based _mt_Ca^2+^ luminescence. The _mt_Ca^2+^ response of *2OX:mtAq* and *mcu*^*1-6*^*:mtAq* mutant seedlings to 400 mM mannitol (a) or 100 mM NaCl (b) treatment is altered compared to Col*:mtAq* wild-type seedlings, respectively. Upper panels are Ca^2+^ luminescence images for demonstrating maximum _mt_Ca^2+^ response, and lower panels are discharge luminescence images for counting total aequorin (see methods for details). FAS imaging was performed with more than six times, and representative images are shown. **c, d** Luminometer reading of Aequorin-based _mt_Ca^2+^ luminescence. The _mt_Ca^2+^ dynamic responses of Col*:mtAq, 2OX:mtAq* and *mcu*^*1-6*^*:mtAq* mutant seedlings to 400 mM mannitol (c) or 100 mM NaCl (d) stimuli. Luminescence intensity was recorded at 1-s intervals, and the arrow indicates the starting point of stimuli application. Each colored line represents the _mt_Ca^2+^ dynamic response from each of 10 individual seedlings. **e, f** Signature of _mt_Ca^2+^ response to mannitol. The _mt_Ca^2+^ peak amplitude (e) and curve area (f) were quantified in 10 individual Col*:mtAq, 2OX:mtAq* and *mcu*^*1-6*^*:mtAq* mutant seedlings treated with 400 mM mannitol, respectively. **g-I** Signature of _mt_Ca^2+^ response to NaCl. The _mt_Ca^2+^ peak amplitude (g), curve area (h) and peak time (i) were quantified in 10 individual Col*:mtAq, 2OX:mtAq* and *mcu*^*1-6*^*:mtAq* mutant seedlings treated with NaCl, respectively. The _mt_Ca^2+^ luminescence peak value was determined by luminometer and converted to _mt_Ca^2+^ concentration as _mt_Ca^2+^ amplitude (see methods for details). The _mt_Ca^2+^ curve area is the normalized luminescence intensity determined by GraphPad Prism 9. a.u. stands for arbitrary unit. **j-k** Cytosolic Ca^2+^ responses are not altered in *2OX:cytAq* and *mcu*^*1-6*^*:cytAq* mutant seedlings. Cytosolic Ca^2+^ response were recorded by FAS imaging, and *2OX:cytAq* and *mcu*^*1-6*^*:cytAq* mutant seedlings show no significant differences from Col*:cytAq* wild-type seedlings in response to 400 mM mannitol (j) or 100 mM NaCl (k) treatment. Luminescence intensity of each FAS film containing about 30 seedlings were determined by indiGO™ software and converted to Ca^2+^ concentration (see methods). Results are expressed as mean ± SEM (*n* = 10 for E-I, n>6 for J and K). Significant differences were examined from one-way ANOVA followed by Dunnett’s multiple comparisons test, \**p* < 0.05, \*\**p* < 0.01. \*\*\**p*<0.001, \*\*\*\**p*<0.0001. See also Supplementary Data 11 for the information of statistical analysis.

### Perturbation of _mt_Ca^2+^ homeostasis triggers mitonuclear protein imbalance and mitochondrial unfolded protein response

To examine whether perturbation of _mt_Ca^2+^ homeostasis may impact mitochondrial function, we analyzed the level of oxidative phosphorylation (OXPHOS) complexes that are encoded by both the mitochondrion (mtDNA) and nuclear (nDNA) genomes in *2OX* and sextuple mutant, together with two other overexpression lines, *MCU4 (4OX)* and *MCU6 (6OX)* (Supplementary Fig. 4). We performed immunoblotting using specific antibodies against subunits of complex l (NADH DEHYDROGENASE2 [NAD2], NAD5, and CARBONIC ANHYDRASE3 [CA3]), lll (APOCYTOCHROME B [COB] and CYTOCHROME C1-1 [CYC1-1]), lV (CYTOCHROME OXIDASE1 [COX1], COX3, and COX X1), and V (ATP SYNTHASE4 [ATP4], ATP8, and ATP2). The abundance of mtDNA encoded COB, COX1, ATP4, and ATP8 are reduced (Fig. 2b-d), but COX3 is increased in *MCU* overexpression and sextuple mutant relative to the wildtype (Fig. 2c), although complex I subunits NAD2 and NAD5 appeared less affected (Fig. 2a). nDNA encoded subunits CA3 and COX X1 both increased (Fig. 2a, c), while CYC1 and ATP2 remained unchanged (Fig. 2b, d). An apparent stoichiometric imbalance between nDNA and mtDNA-encoded subunits was evident for complexes lll, lV, and V (Fig. 2b-d).

**Fig. 2.**
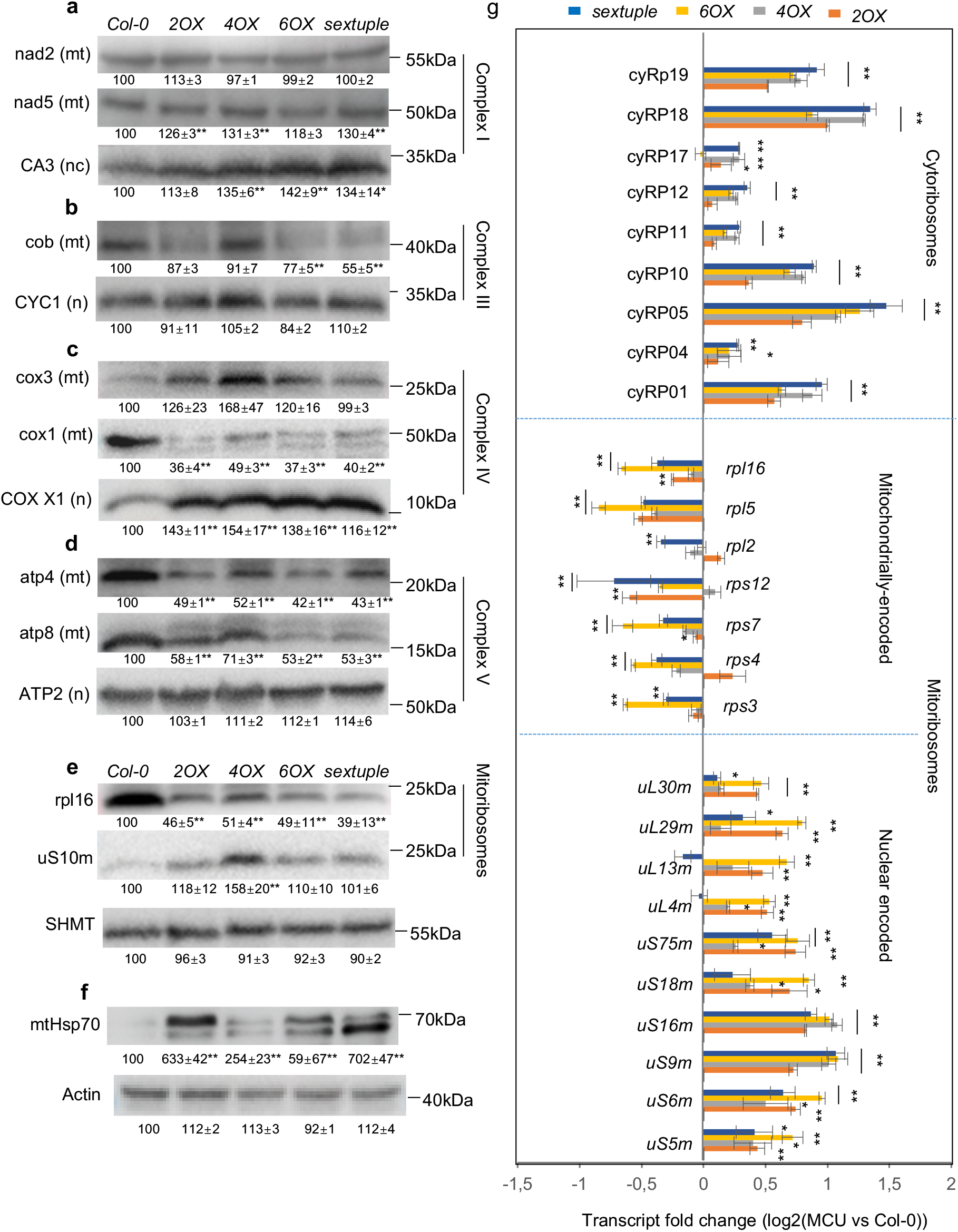
Abundance of OXPHOS subunits is altered in both gain-of-function and loss-of-function *MCU* plants. **a-d** Comparison of OXPHOS subunit protein abundance. nDNA subunits (labeled as n) and mtDNA subunits (labeled as mt) of OXPHOS complexes I (A), lll (b), lV (c), and V (d) were detected and quantified by immunoblotting using antibodies against each subunit. (e) Comparison of ribosome proteins. Abundance of mtDNA rpl16 proteins and nDNA uS10m were altered in *MCU* plants. **f** Comparison of mitochondrial chaperone. nDNA mitochondrial chaperone mtHsp70 was detected using specific antibody. SHMT and actin proteins are loading controls for mitochondrial protein and cytosolic proteins, respectively. Quantifications of signals relative to the wild type (set to 100%) are provided below each panel as means ± SEM (*n* = 3). Two-tailed student’s *t*-test was performed to examine statistical significance, \**p* < 0.05, \*\**p* < 0.01 with fold change (vs wildtype) |FC|>1.2. **g** Relative transcript levels of mitoribosome (*mtRP*) and cytoribosome (*cyRP*) genes encoded by nDNA or mtDNA, as determined by RT-qPCR. The results are expressed as the Log_2_ fold-change in *MCU* overexpression and sextuple mutant relative to wild-type and shown as means ± SEM (*n* = 3). Two-tailed student’s *t*-test was performed to examine statistical significance, \**p* < 0.05, \*\**p* < 0.01. See also Data10 for the information of antibodies and primers, and Supplementary Data 11 for the information of statistical analysis.

Next, we examined the transcript levels of mtDNA encoded mitochondrial ribosomes (*rps*) and nDNA encoded mitochondrial ribosomes (*mtPRs*) and cytosolic ribosomes (*cyRP*s) using RT-qPCR. Most mtDNA *mtRP* transcripts were downregulated, while nDNA *mtRP* and *cyRP* transcripts were upregulated in *MCU* overexpression and sextuple mutant plants compared to the wildtype, suggesting that nDNA and mtDNA gene expression is not coordinated at the transcriptional level (Fig. 2g). Using specific antibodies, we observed that the abundance of the mtDNA rpl16 decreased while nDNA uS10m increased in *MCU* overexpression and sextuple mutant plants (Fig. 2e). Moreover, the abundance of mitochondrial chaperone protein mtHsp70 is higher in *MCU* overexpression and sextuple mutant plants relative to wildtype (Fig. 2f). Together, these results suggest that perturbation of MCU-controlled _mt_Ca^2+^ homeostasis causes the misregulation of mtDNA and nDNA ribosome gene expression. Activation of UPR^mt^ implies that perturbation of _mt_Ca^2+^ homeostasis initiates a retrograde communication with the nucleus.

### Impaired MCU controlled _mt_Ca^2+^ homeostasis activates a multi-compartment proteostatic response

To assess whether **i**mpaired **M**C**U** controlled _mt_**C**a^2+^ **H**omeostasis (iMUCH) may trigger an undefined retrograde signaling pathway communicating with nuclei, we performed RNA-seq analysis of *2OX and sextuple* mutant together with wildtype grown under both normal and mitochondrial stress conditions (exposure to the electron transport chain inhibitor antimycin A [AA]). We identified 2,458 and 954 differentially expressed genes (DEGs) in *2OX* and sextuple mutant respectively relative to the wildtype under normal conditions. To our surprise, among the total DEGs, genes in the categories ‘ribosome biogenesis and translation’, ‘cytosolic ribosome’ and ‘mitochondrial ribosome’, were the top enriched GO biological function and cellular component terms, respectively (Supplementary Data 1). We examined the distribution of transcripts and found that most of the translation, cytosolic translation, cytosolic ribosome, and mitochondrial ribosome genes were upregulated in both *2OX* and sextuple mutant (Fig. 3a and Supplementary Data 2). Moreover, in addition to mtHsp70, many more nDNA encoded mitochondrial chaperone genes were significantly induced in both *2OX* and sextuple mutant compared to wildtype (Fig. 3b). We further examined the expression of 21 genes encoding cytosolic chaperones (cyHsp), 11 genes encoding endoplasmic reticulum (ER) chaperones (erHsp), as well as 14 genes encoding chloroplast chaperones (cpHsp). These chaperone genes were also transcriptionally induced in *2OX* and sextuple mutant compared to wildtype (Fig. 3b; Supplementary Data 3). It is known that AA induces mitochondrial stress and triggers mitochondrial retrograde response (MRR) ^43-47^. However, neither the cytosolic nor the mitochondrial ribosome genes were enriched in the DEGs identified in AA-treated wildtype or AA-treated *2OX* and sextuple mutant (Supplementary Data 4). In addition, the expression of chaperone genes was not affected by AA treatment, with the exception of ER and chloroplast chaperone genes, whose transcript levels decreased slightly in AA-treated *2OX* and sextuple mutant, but not in AA-treated wildtype (Supplementary Fig. S4 and Supplementary Data 3).

**Fig. 3.**
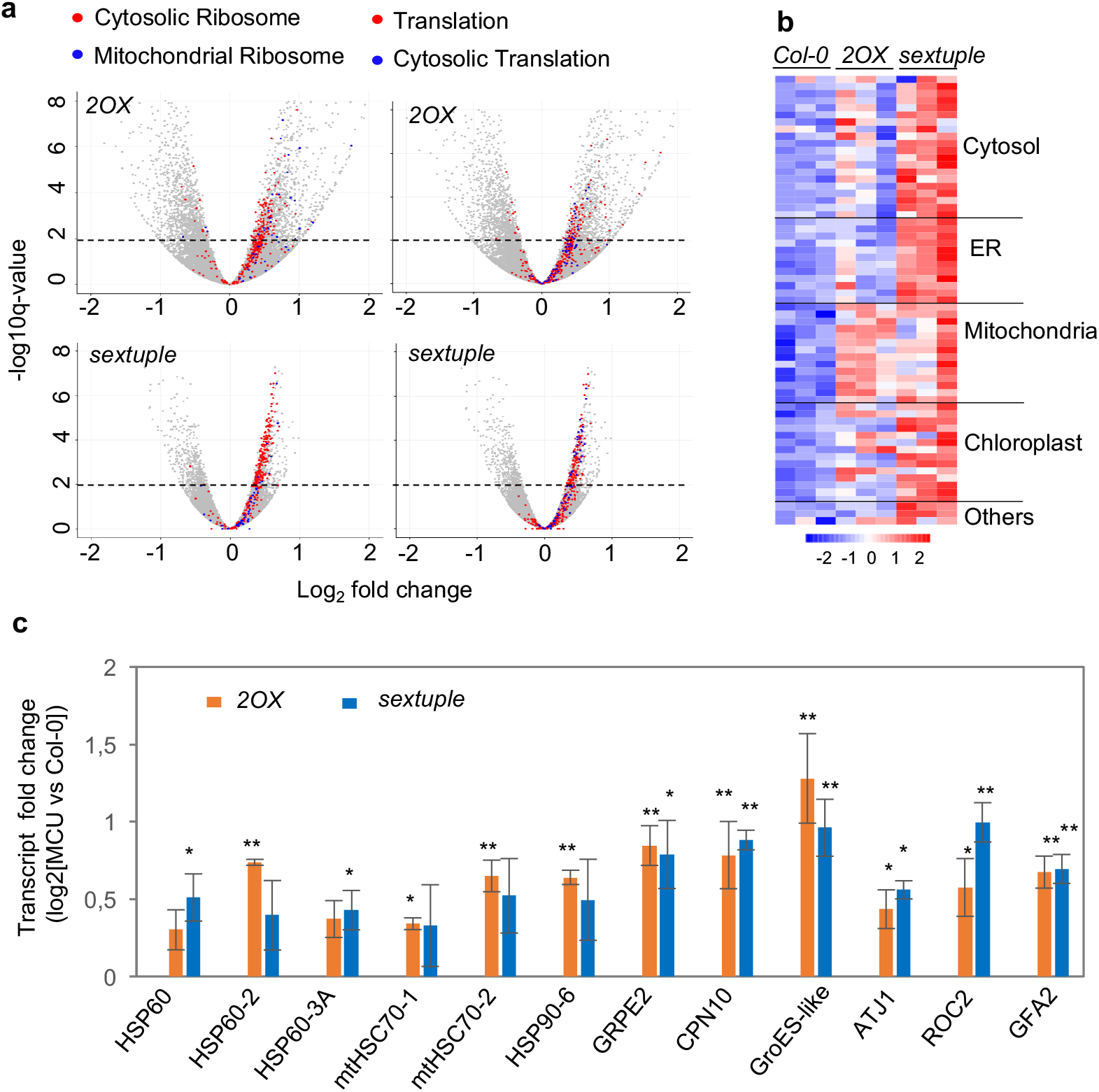
iMUCH induces nDNA *RPs* and multi-compartment chaperone gene expression. **a** Distribution of differentially expressed transcripts quantified by RNA-seq in *2OX* and *sextuple* mutant relative to wildtype plants, shown as Volcano plots, highlighting GO terms linked to cytosolic ribosome and mitochondrial ribosome (left panel) and translation and cytosolic translation (right panel) as provided by the Arabidopsis plant genome database (http://plantgdb.org/AtGDB/). See also Supplementary Data 4. **b** Induction of multi-compartment UPR gene expression. Heatmap representation of chaperone gene expression, indicating differentially expressed chaperone genes of the cytosol, the ER, the mitochondria, and the chloroplast in *2OX* and *sextuple* plants. See also Supplementary Data 3 for the list of gene ID. **c** Induction of mitochondrion-localized chaperone gene expression. Relative transcript levels of mitochondrial chaperone genes were quantified by RNA-seq, highlighting that mitochondrial chaperone genes are highly induced in *2OX* and *sextuple* mutant compared to wildtype plants. Results are expressed as means ± STDEVP (n=3). Two-tailed student’s *t*-test was performed to examine statistical significance, \**p* < 0.05, \*\**p* < 0.01. See also Supplementary Data 11 for the information of gene ID and statistical analysis.

Our analysis identified 2,384 AA-induced DEGs overlapping across wildtype, *2OX* and *sextuple* (adjusted *p*-value < 0.05) (Supplementary Fig. 6A and Supplementary Data 4). We identified 199 genes encoding transcriptional regulators (GO:0140110) among overlapping DEGs (Supplementary Data 5), including mediators of retrograde signaling, NAC013 and NAC053 that regulate the expression of *ALTERNATIVE OXIDASE 1a* (*AOX1a*) ^48^. WRKY40 and WRKY63 that regulate genes encoding mitochondrial proteins ^49^. TF genes that displayed an AA-induced up- or down-regulation in all samples are strong candidate mediators of retrograde signaling (Supplementary Data 5). Interestingly, fold-changes in MRR marker genes were much higher in *2OX* and sextuple mutant compared to wildtype (Supplementary Fig. 6B). Moreover, the expression levels of AA-induced TF genes were two-fold higher in *2OX* and sextuple mutant than in the wildtype, including *NAC032* and *NAC102, WRKY45* and *WRKY75, ZAT8* and *ZAT12, DEHYDRATION-RESPONSIVE ELEMENT BINDING PROTEIN 2A* (*DREB2A*) and *DREB2C, SCL1* and *SCL13*, and *HSFA4A* and *HSFA3* (Supplementary Fig. 5C). These analyses revealed that both gain-of-function and loss-of-function *MCU* activates a multi-compartment *UPR* and *RP* gene expression, and general stress response, highlighting that mitochondrial cross-compartmental signaling is imitated likely to maintain cellular proteostasis upon iMUCH.

### Selective translational repression of RNA modification and ribosomal gene expression are evident in both gain-of-function and loss-of-function *MCU* plants

Transcriptional activation of cytosolic *UPR* and *RP* genes imply that a cytosolic proteostasis response may occur at the translational level upon iMUCH. To test this possibility, we performed polysome profiling in *2OX*, sextuple, and wildtype to assess the status of cytosolic translation ^50^. We separated free ribosomal subunits, monosomes, and polysomes over a sucrose density gradient, followed by quantification by optical density. We observed a shift from polysomes to monosomes in *2OX* and sextuple mutant plants relative to the wildtype, suggestive of a global translational decrease (Fig. 4a, upper panel). To establish whether this response was due to iMUCH, we also performed the polysome profiling in AA-treated wildtype. AA treatment caused a significant shift from polysomes to monosomes (Fig. 4a, lower panel). These observations suggest that iMUCH and impairing ETC function cause a global translational repression.

**Fig. 4.**
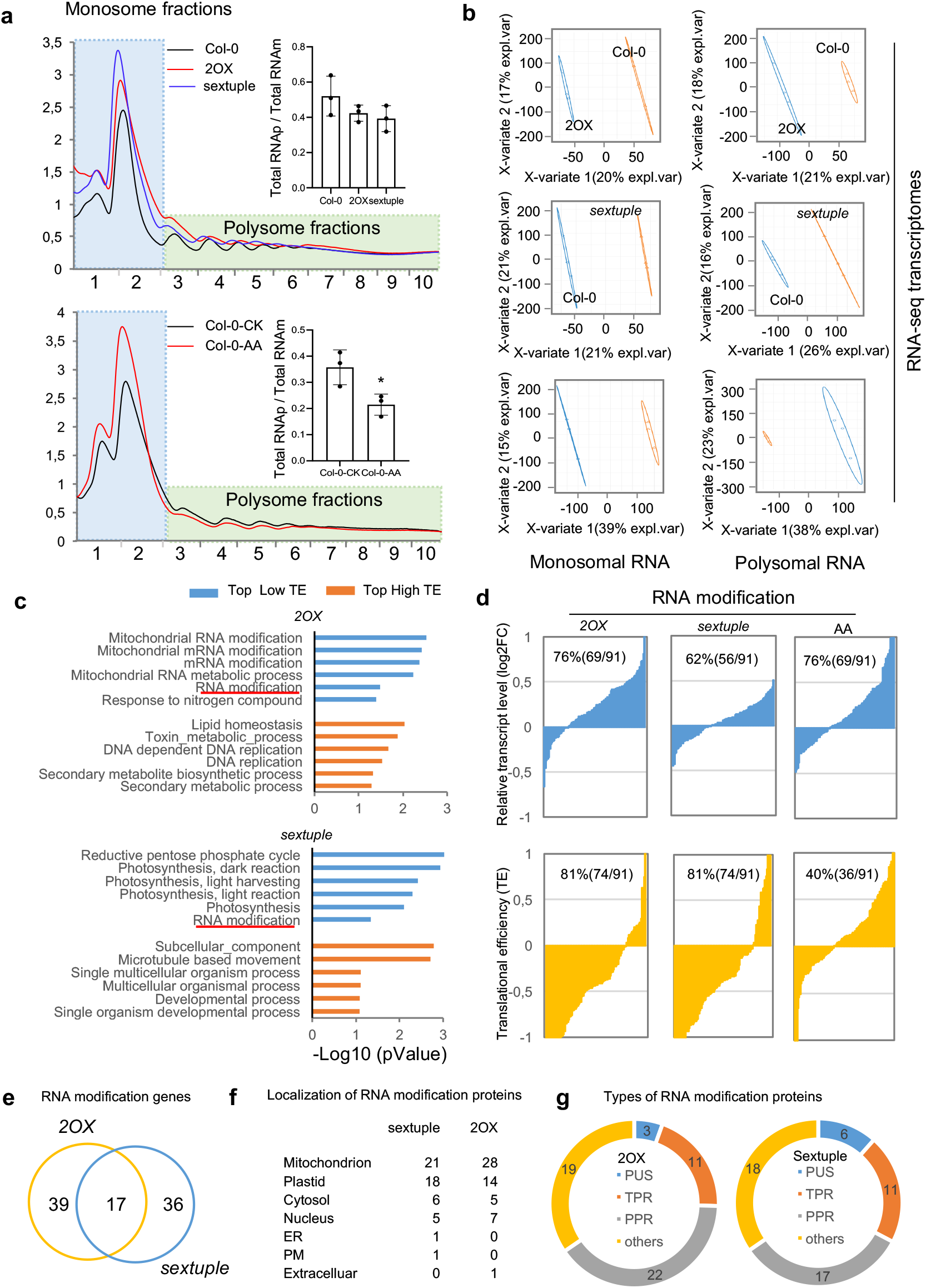
*2OX* and *sextuple* mutant display reduced translational efficiency of RNA modification proteins. **a** Representative polysome profiles showing altered cytosolic polysome abundance in *2OX* (red line) and sextuple mutant (blue line) relative to the wildtype (black line) (upper panel) and in AA-treated (red line) and control (black line) wildtype (lower panel). Monosomal fractions and polysomal fractions are indicated in the blue and green backgrounds, respectively. RNAs isolated from highly translated fractions (green, polysomes) and lesser translated fractions (blue, monosomes) were used for RNA-seq analysis for *2OX*, sextuple mutant, and wild-type as well as AA-treated and control wildtype. Inserted panels are the histogram of total polysomal RNA (RNAp) divided by total monosomal RNA of 2OX, sextuple mutant and wildtype (upper insertion), and that of AA-treated and untreated wildtype (lower insertion). **b** Partial least-squares discriminant analysis (PLS-DA) for monosomal (left) and polysomal (right) RNA-seq libraries showing the distinction between *2OX* (upper panel), *sextuple* mutant (middle panel) and wildtype controls, and between AA-treated and untreated wild-type (lower panel). **c** Top six biological processes with highest enrichment scores for low translational efficiency (TE, blue) or high TE (yellow) transcripts in GO. TE is defined as the Log_2_ ratio of polysomal versus monosomal differences on transcript levels between *2OX, sextuple* mutant and control wildtype or AA-treated and untreated wild-type. Clusters with an “enrichment score” above 1 (*p* < 0.001) were considered as significant ones. **d** Visualization of transcript and TE levels of mRNAs enriched in RNA modification. Proportion of downregulated and upregulated transcripts (upper panel) and their TEs (lower panel) were plotted for *2OX*, sextuple mutant, and AA-treated wildtype relative to their controls. The percentage of upregulated transcripts and transcripts with low TEs is indicated within each plot. **e-g** Identity of genes enriched in RNA modification. The Venn diagram shows the number of overlapping genes and unique genes detected in *2OX* and sextuple mutant, respectively (e), with their predicted subcellular localization (f), and the number of genes encoding PPR-related proteins in both *2OX* and sextuple mutant (g).

Having observed the decreased global translation in *2OX* and sextuple mutant, and in AA-treated wildtype, we next determined which genes are repressed translationally. We generated RNA-seq libraries of the monosomal and polysomal RNA fractions. This RNA-seq dataset provided information for over 23,000 genes, covering the majority of the Arabidopsis genome and allowing for a global description of the differences between monosomal and polysomal fractions (Supplementary Data 6). We performed a partial least-squares discriminant analysis (PLS-DA) to distinguish *2OX* and the sextuple mutant from wildtype, and AA-treated and untreated wildtype (Fig. 4b). Next, we calculated the translational efficiency (TE) of each gene (see Methods). A positive Log_2_ TE value indicates a higher TE for a given transcript that shifts from monosomes to polysomes, whereas a negative Log_2_ TE value represents a lower TE of a given transcript that shifts from polysomes to monosomes. We applied a minimal fold-change TE cutoff of 1.25 with a significant Variable Importance in Projection (VIP) score to filter genes with significantly altered TE (saTE) for GO term clustering (Supplementary Data 6). Genes with a high saTE encoded proteins involved in lipid homeostasis and secondary metabolism in *2OX*, and in protein transport and establishment of protein localization in sextuple mutant. This observation likely reflects the activation of metabolic adaptation and the enhancement in inter-organellar contacts ^8,51^. Photosynthesis related genes with low saTE were highly enriched in sextuple mutant, implying that the translation of most photosynthesis related genes is selectively repressed in sextuple mutant. Intriguingly, genes with low saTE in both *2OX* and sextuple mutant, but not in AA-treated wildtype, are enriched in mitochondrial RNA metabolism and RNA modification (Fig. 4c and Supplementary Fig. 7), indicating that the selective translational repression of RNA modification genes takes place upon iMUCH. We next inspected our RNA-seq data and determined that about 76%, 62%, and 65% of RNA modification genes tend to be upregulated in *2OX*, sextuple mutant, and AA-treated wildtype, respectively. About 81% of their transcripts with low TE were detected in *2OX* and sextuple mutant, but only 41% in AA-treated wildtype, indicating again the high translational suppression of RNA modification genes in *2OX* and sextuple mutant (Fig. 4d). Among the enriched RNA modification genes with low saTE, we identified 56 (*2OX*) and 53 (sextuple mutant) genes as significant hits, of which 17 genes were shared by *2OX* and sextuple mutant (Fig. 4e). Of those, about half encoded mitochondrion- or chloroplast-localized PPR, TPR, and PUS proteins, which regulate organellar functions by executing organellar RNA editing and translation (Fig. 4f, g). Genes with saTE were enriched mostly in organic acid metabolism and response to oxygen level in AA-treated wildtype (Supplementary Fig. S7), indicating that translational adaptation mostly works towards metabolism and stress response to AA-induced mitochondrial stress.

Although *cyRP*s were not part of the top list of enriched genes with saTE, we also assessed their translational efficiency. About 88% and 93% of *cyRPs* were transcriptionally upregulated in *2OX* and sextuple mutant, respectively (Supplementary Fig. 8A and Supplementary Data 6). However, we failed to detect a similar transcriptional regulation of *cyRPs* in AA-treated wildtype, in which about half (52%) of *cyRPs* tended to be upregulated upon AA treatment (Supplementary Fig. 8A and Supplementary Data 6). A large proportion of *cyRP*s had low TE in *2OX* (76% of *cyRP*s) and sextuple mutant (62% of *cyRP*s), and 51% of *cyRP*s with low TE were detected in AA-treated wildtype, implying a strong repression of cytosolic translation of *cyRP* genes in *2OX* and sextuple mutant (Supplementary Fig. 8B and Supplementary Data 6). We independently validated the translational repression of some *cyRP*s and *mtRP*s by RT-qPCR in *2OX* and sextuple mutant. In addition to nDNA encoded *cyRPs and mtRPs*, mtDNA encoded *mtRPs* were repressed translationally (Supplementary Fig. 8C, D). These findings suggest that genes in organellar and cytosolic translational machinery are subjected to translational regulation upon iMUCH. Although the global translational response occurred in *2OX* and sextuple mutant, and AA-treated wildtype with various degrees of repression, a prominent translational repression of RNA modification and cytosolic ribosome proteins took place in 2*OX* and sextuple mutant, but not in AA-treated wildtype. These results clearly demonstrate that iMUCH, but not AA triggers a cytosolic translation response with specific impacts on RNA modification protein and cytosolic ribosome protein synthesis.

### Post-transcriptional repression of ribosomal gene expression is the molecular consequence of IMUCH

To better understand how iMUCH evoked translational response determines the proteome, we performed a mass spectrometry analysis of the proteome in the sextuple mutant and the wildtype. To identify the proteome signatures of the sextuple mutant, we analyzed the proteome of AA-treated wildtype in parallel. Proteomics quantified over 6,000 proteins in each sample (Supplementary Data 7). With filtering criteria of *p*-value < 0.05 and |Log_2_FC| > 0.263 (|FC|>1.2), we identified 2,240 and 116 differentially abundant proteins (DAPs) in the sextuple mutant and in AA-treated wildtype, respectively. To examine how the changes in transcript and protein levels were correlated in cellular process, we performed a co-regulation analysis by plotting the fold-change (FC) of DAPs versus the FC of transcript levels for each individual gene and protein (Fig. 5a) for 2,212 protein–transcript pairs with the above criteria applied to proteins and no criteria applied to transcripts (Supplementary Data 8). Our co-regulation analysis revealed that about half of all genes were coordinately regulated, with the same direction of regulation in transcript and protein levels, while the remaining half of genes were regulated differently at the protein and transcript levels in sextuple mutant (Fig. 5a). Indeed, 24% of all genes were co-upregulated (Q1) and 28% were co-downregulated (Q3), with a relatively smaller portion (15%) of genes being downregulated at the transcript level and upregulated at the protein level (Q2). However, a relatively larger portion of genes (33%) exhibited an upregulation at the transcript level and a downregulation at the protein level (Q4) (Supplementary Data 8).

**Fig. 5.**
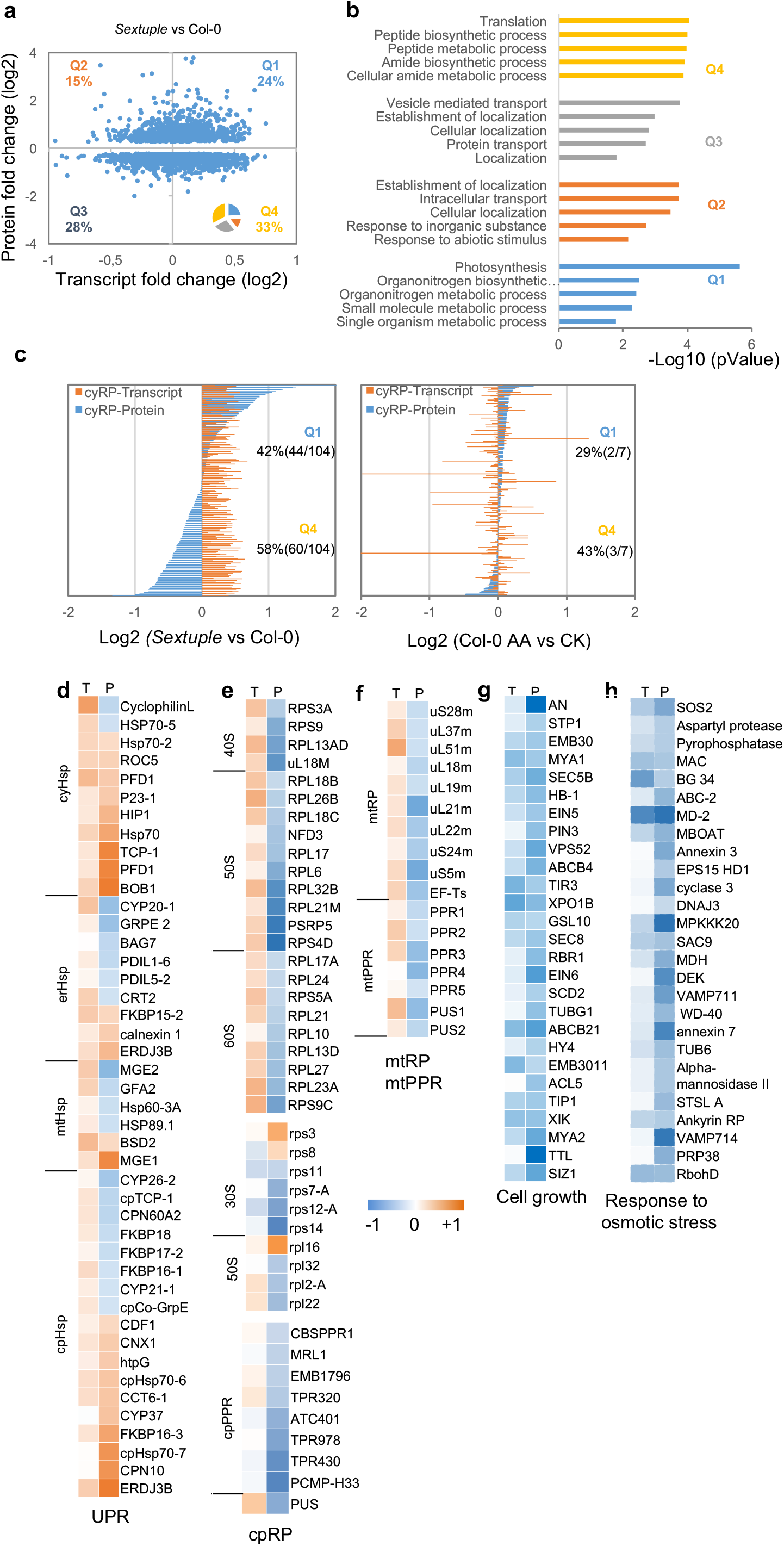
Proteome–transcriptome cross analysis reveals post-transcriptional repression of ribosome gene expression and transcriptional and post-transcriptional co-repression of growth and stress response pathway. **a** Co-regulation plot of transcript and protein fold-change (FC) of sextuple mutant versus wildtype plants. The percentage of co-upregulated proteins (Q1), post-transcriptionally increased proteins (Q2), co-downregulated proteins (Q3), and post-transcriptionally repressed proteins (Q4) is indicated in each quadrant. **b** Top five biological processes enriched in each quadrant of the co-regulation plot, derived from the significant hits of differentially abundant proteins (DAPs). See also Supplementary Data 8. **c** Co-regulation of cyRPs. Individual transcripts (yellow) and proteins (blue) were plotted with Log_2_(FC) of sextuple mutant versus wildtype plants (left panel) and AA-treated wildtype versus untreated control (right panel), demonstrating a large proportion of post-transcriptionally repressed cyRPs in sextuple mutant, but not in AA-treated wildtype. **d-h** Examples of co-regulated proteins identified by mass-spectrometry. Most cyHSPs (9 out of 11) and proposition of erHSPs (3 out of 9), mtHSPs (2 out of 6), and cpHsp (10 out 18) are co-upregulated (d). Examples of most cpRPs and cpPPRs (e) and mtRPs and mtPPRs (f) in Q4 that are transcriptionally induced, but post-transcriptionally repressed. An example of co-downregulated proteins in cell growth (g) and response to osmotic stress (h) is also demonstrated. T: transcript, P: protein. See also Supplementary Data 8 for the list of gene ID.

To pinpoint the biological function of genes within each Q1–Q4 quadrant, we performed GO enrichment analyses for their encoded proteins. We showed the top five terms based on their *p*-value and ordered them as a function of their enrichment score (Fig. 5b and Supplementary Data 8). Genes with upregulated transcripts and downregulated proteins (Q4) showed enrichments for translation and peptide biosynthetic processes. Manual inspection of *RP* genes involved in cytosolic translation revealed that 104 out of 194 cyRPs are detected as DAPs, of which 42% belong to Q1 and 58% to Q4, indicating that about half of all *cyRP* genes are repressed post-transcriptionally (Fig. 5c and Supplementary Data 8). Only seven cyRPs were detected as DAPs in AA-treated wildtype, with two in the Q1 and three in the Q4 quadrant (Fig. 5c and Supplementary Data 8). Upon manual inspection, we identified 23 nDNA chloroplast ribosome proteins (*cpRP*s), 5 chloroplast-encoded (cpDNA) *cpRP*s (Fig. 5e), and 9 nDNA *mtRP*s (Fig. 5f) as well as a group of genes encoding mitochondrion- and chloroplast-localized pentatricopeptide repeat proteins (PPRs), tetratricopeptide repeat proteins (TPRs), and pseudouridine synthases (PUSs) involved in organellar RNA editing and translation in Q4 (Fig. 5e, f). In addition, we detected 11 DAPs in translation initiation (GO:0006413), including two eIF-3 (AT4G20980 and AT5G44320) that function in cap-dependent translation initiation (GO:002191), and two GCN2 targets, eIF2α (AT2G40290 and AT5G05470), and 10 DAPs in ribosome assembly (GO:0042255) in Q4 (Supplementary Data 8). These results indicate that a large proportion of cytosolic and organellar ribosome proteins, as well as proteins with roles in cytosolic and organellar RNA editing and translation, are subjected to post-transcriptional repression, which is in line with TE analysis. Organonitrogen metabolism and photosynthesis were ranked at the top of co-upregulated gene enrichment terms for Q1, which likely reflects a part of active adaptation to disturbed mitochondrial function. Manual inspection of Q1, we found 18 DAPs in protein folding (GO:000645) that are cytosolic-, mitochondrial-, chloroplast- and ER-localized chaperones (Fig. 5d). The enrichments in “vesicle-mediated transport” and “auxin transport” in Q3 indicate the reduction of cell wall biogenesis and hormone controlled growth. Again, we identified genes encoding proteins for cell growth (GO:0016049) and response to osmotic stress (GO:0006970) (Fig. 5g, h). Together with development, and maturation as significant hits with enrichment score >1 in Q3 (Supplementary Data 8), this suggests a transcriptional and translational co-repression of growth related genes and stress responsive genes upon iMUCH. Together, this iMUCH evoked post-transcriptional response highlights mitochondrial stress dependent regulation of protein synthesis with impacts on multi-compartmental proteostasis.

### GCN2-eIF2α but not TOR pathway is involved in iMUCH-induced translational regulation

Protein synthesis can be attenuated by the decrease of mTOR signaling activity or the increase in the phosphorylation of eukaryotic translation initiation factor 2 (eIF2α ^52,53^. To identify the potential effectors, we first analyzed the expression of components in the TOR-S6 kinase (S6K) and GENERAL CONTROL NON-DEPRESSIBLE2 (GCN2)-eIF2α pathways, and detected that *TOR, REGULATORY-ASSOCIATED PROTEIN OF TOR1* (*RAPTOR1*), *S6K2, GCN2, ATP-BINDING CASSETTE F3* (*ABCF3*), *ABCF4*, and *ABCF5* were expressed to lower levels, while *RIBOSOMAL PROTEIN S6A* (*RPS6A*), *E2F TRANSCRIPTION FACTOR3* (*E2F3*), *ERBB-3 BINDING PROTEIN1* (*EBP1*), *eIF2*α, and *eIF2-A2* were more highly expressed in *2OX*, sextuple mutant, and AA-treated wildtype (Fig. 6a, b). To check for the transcriptional and translational co-regulation of these key components, we mined the proteomic data and discovered that eIF2α, eIF2-A2, ABCF4, and ABCF5 protein levels are significantly lower in the sextuple mutant, although most TOR components were not detected by mass spectrometry. We observed a similar trend for eIF2α and eIF2-A2, but their downregulation did not reach significance in AA-treated wildtype (Fig. 6g). Besides, other components in translation initiation also showed a significant reduction, including eIF3d (cytoplasmic cap-dependent translation initiation) and EIF3, TRM61 and NagB (regulation of translation initiation) (Fig. 6d). To substantiate these observations, we further examined the abundance of p-eIF2α (phosphorylated form of eIF2α), KIN10, and the phosphorylated form of KIN10 (p-KIN10), KIN11 (p-KIN11), and TOR protein by immunoblotting. The p-eIF2α levels were significantly reduced, but p-KIN10 and TOR abundance showed no obvious changes in *2OX* or sextuple mutant although a slight increase of p-KIN11 in *2OX* and decrease of KIN10 in sextuple mutant were detected (Fig. 6c, e). By contrast, we detected a significant increase of p-KIN10 and p-KIN11 and reduction of TOR, but no obvious alteration of p-eIF2α abundance in AA-treated wildtype, although KIN10 showed no changes in AA-treated wildtype (Fig. 6d, f). These results suggest that the translational response to iMUCH and AA-induced mitochondrial stress may be transduced by different pathways.

**Fig. 6.**
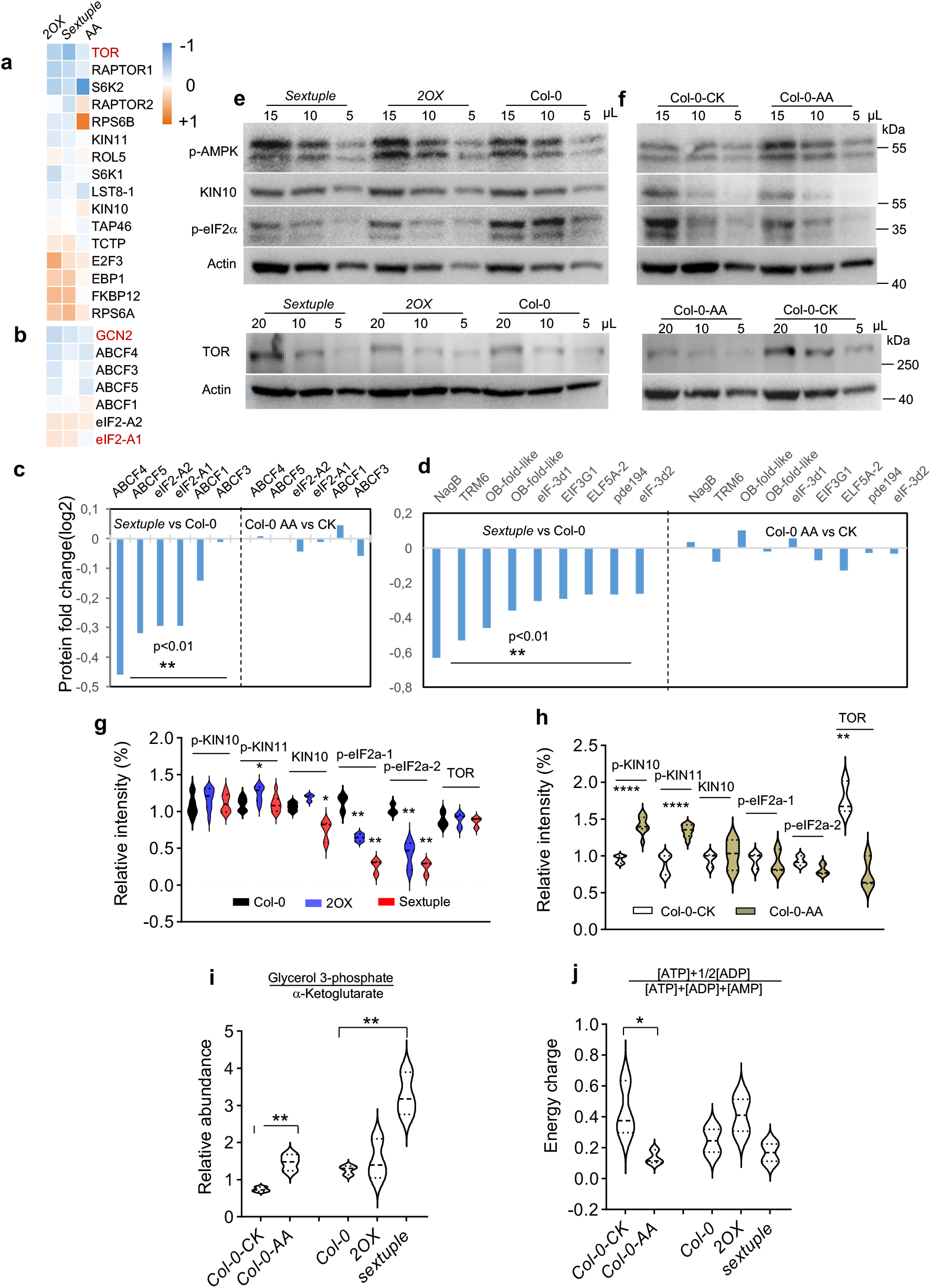
The GCN2-eIF2α but not TOR-S6K pathways respond to iMUCH. **a, b** Heatmap representation of the transcriptional regulation of the components in the TOR-S6K (a) and GCN2-eIF2α pathways (b), based on Log_2_(FC) between the *2OX* versus Col-0, between *sextuple* mutant versus Col-0, and between AA treated Col-0 versus untreated controls, as indicated as 2OX, sextuple, AA on the top of the heatmap panel. **c, d** Protein abundance of components in GCN2-eIF2α pathway (c) and in translation initiation (GO:0006413) (d). Comparison of protein abundances between the sextuple mutant and Col-0, and between AA treated Col-0 versus untreated controls was performed by quantitative proteomic analysis. Significant hits are those with |Log_2_(FC)| > 0.263 and *p* < 0.01. **e, f** Protein abundances of the components in TOR and eIF2α pathway were determined by immunoblotting. Samples and their loading volumes are indicated at the top of the panel. Actin serves as loading control. Anti-p-AMPK antibody recognizes p-KIN10 (upper band) and p-KIN11 (lower band). **g, h** Quantification of protein abundance. Signal intensity was quantified with at least three immunoblots using Evolution-Capt software. Significant differences were examined with one-way ANOVA followed by Dunnett’s multiple comparisons test for e, \*\**p* < 0.01. \*\*\*\**p*<0.0001. Two-tailed student *t*-tests were performed for f, \*\*\*\**p*<0.0001. **i** Metabolic shifting. The ratio of the relative abundance between glycerol 3-phosphate (glycolysis) and α-ketoglutarate (TCA cycle) was calculated for the *sextuple* mutant and wildtype, as well as for AA-treated and untreated wildtype. A similar metabolic shift toward glycolysis was observed in *sextuple* and AA-treated wildtype, respectively. Significant differences were examined with two-tailed student’s *t*-tests (\*\**p* < 0.01, *n* = 3). **j** Energy status. Energy charge was measured as the index of ATP, ADP, and AMP. Significant differences (P < 0.05) were examined with two-tailed student’s *t*-tests (\**p* < 0.05, *n* > 2). See also Supplementary Data 11 for the information of statistical analysis.

To further dissect the pathways that mediate the translational response to iMUCH, we determined energy-related metabolites. We detected reduced contents for tricarboxylic acid (TCA) cycle intermediates (α-ketoglutarate, citrate, succinate, and oxaloacetate) and higher levels of glycolysis intermediates (3-phosphoglycerate and lactate) in *2OX* and sextuple mutant compared to wildtype (Supplementary Data 9). Calculating the ratio between α-ketoglutarate and 3-phosphoglycerate, we noticed a shift from TCA metabolism toward glycolysis in 2OX, sextuple mutant and AA-treated wildtype, although it is not significant in 2OX plants (Fig. 6h). We also estimated the energy charge (ENC), an index used to measure the energy status. We obtained a lower ENC in AA-treated wildtype but not in *2OX* and sextuple mutant (Fig. 6i), suggesting that iMUCH alters mitochondrial metabolism but has no significant effects on cellular energy status. Thus, the GCN2-eIF2α and TOR-S6K pathways mediate the cytosolic translational response to mitochondrial stress with the TOR-S6K pathway sensing cellular energy status and the GCN2-eIF2α responding to iMUCH in an energy-independent manner.

### iMUCH is coupled with the reduction of growth and stress resistance

Connection between mitochondrial and cytosolic proteostasis is likely related to organismal health and cellular stress resistance ^54-56^. To determine the physiological consequence of iMUCH evoked proteostatic response, we thus examined the growth and stress resistance of sextuple mutant and four independent *2OX* transgenic lines. We first tracked the seedling growth and development of four *2OX* lines. It appeared that the size of *2OX* plants is correlated with the level of MCU2 (Fig. 7c, d), indicating that the growth inhibition is related to MCU2 expression. We compared the size of 20-day-old seedlings by measuring leaf size and number, and found a significant reduction in the leaf area but not in leaf number of *2OX* and sextuple mutant compared to wildtype (Fig. 7a, b). Differences in growth were minimized as the developmental process in sextuple mutant. In addition, *2OX* lines and sextuple mutant showed leaf premature senescence under normal growth conditions. The timing of leaf premature senescence also appeared to be *MCU2* level-dependent (Fig. 7c, d). However, as the plants matured, all *2OX* lines showed earlier leaf senescence compared to the wildtype (Fig. 7c). Sextuple mutant exhibited leaf premature senescence in late stage under normal growth conditions (Fig. 7d). We tested the resistance to mannitol induced osmotic stress, and found that the growth of *2OX* and sextuple mutant was significantly inhibited compared to the wildtype (Fig. 7f, g). These results show that iMUCH shapes plant growth, development and stress resistance.

**Fig. 7.**
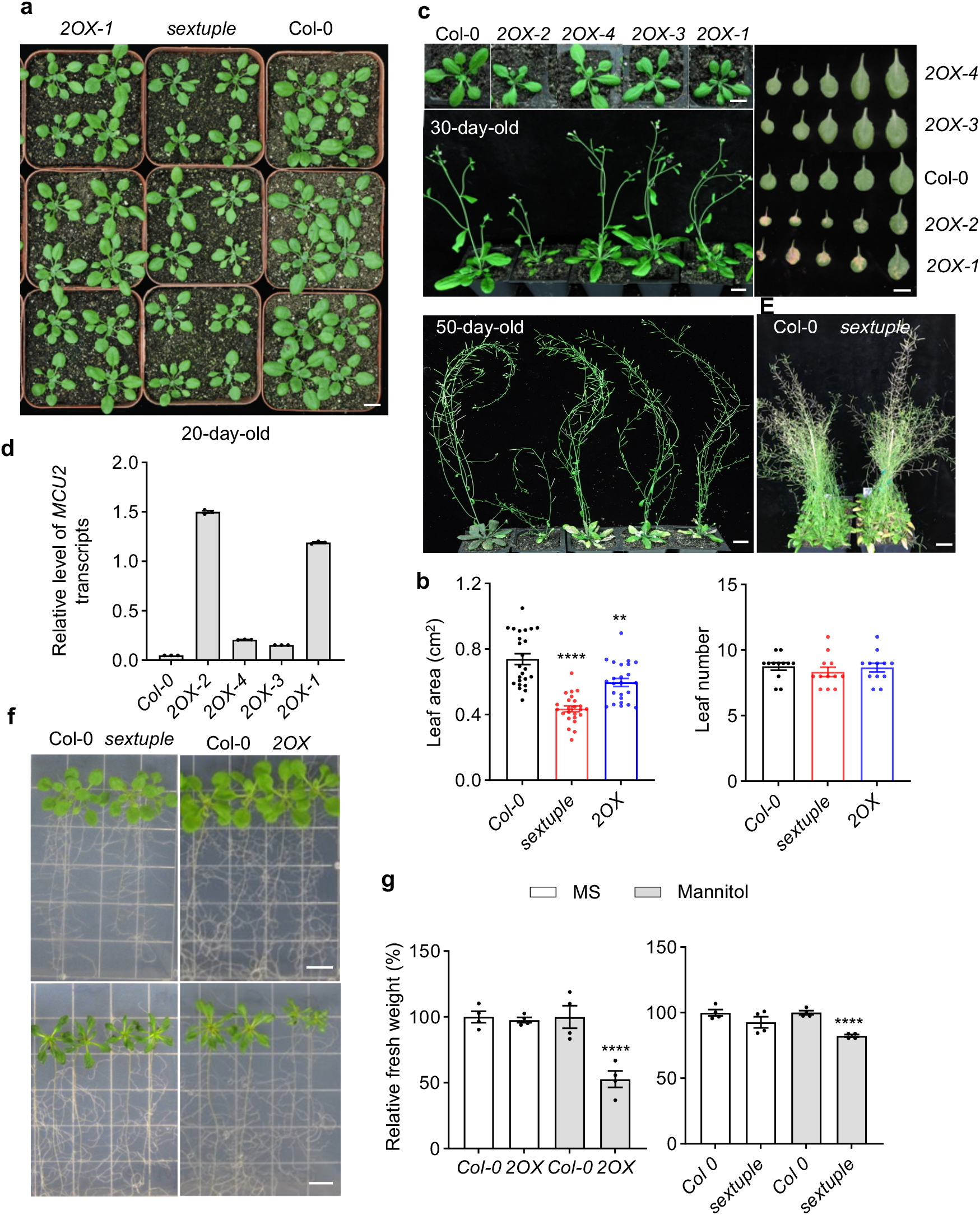
*2OX* and *sextuple* mutant show reduction in growth and osmotic stress resistance and acceleration of leaf senescence. **a, b** Morphological comparison of seedlings grown in soil under normal growth condition. Morphology of 10-day-seedlings (a), the area of two largest rosette leaves and the number of leaf of 10-day-old seedlings (b) were recorded. Results are shown as the mean ± SEM (n=24). Two-tailed student’s t-tests were performed. \*\*\**p*<0.001, \*\*\*\**p*<0.0001. **c** *2OX* plants show MCU2 expression level-dependent early leaf senescence. **d** Relative transcript levels in *MCU2* transgenic lines measured by RT-qPCR. **e** Sextuple mutant show accelerated leaf senescence compared to wild-type plants at the mature stage. **f** 2OX and sextuple mutant are sensitive to mannitol-induced osmotic stress. Five-day-old seedlings were transferred to plates containing MS medium only (upper panel) or MS medium containing 150 mM mannitol (lower panel). Photographs were taken 10 days after transfer. **g** Relative fresh weight of *2OX* and sextuple seedlings compared to wild type, measured 10 days after transfer. Results are shown as the mean ± SEM (n=4), normalized to wild-type values (set to 100%). Significant different levels were examined with two-tailed student’s t-tests. \*\*\*\**p*<0.0001. See also Supplementary Data 11 for the information of statistical analysis. Scale bar is 1 cm.

## Discussion

In this study, we demonstrate that mitochondrion-localized Ca^2+^ uniporters are necessary for _mt_Ca^2+^ homeostasis *in planta*. Although both Arabidopsis *MCU1* and *MCU2* are sufficient to generate mitochondrial Ca^2+^ influx when expressed in yeast, their roles in controlling _mt_Ca^2+^ responses *in planta* had not been established ^34,35^. A recent study reported the significantly reduced _mt_Ca^2+^ basal level and the mitochondrial Ca^2+^ uptake in the root cell of Arabidopsis *mcu1 2 3* triple mutant^38^, which matches the expression pattern of *MCU1, MCU2 and MCU3* in root cells near the root tip. And the residual Ca^2+^ uptake activity in the *mcu1 2 3* line is likely due to the predominant expression of MCU6 in roots (Supplementary Fig. S1C). Because of significantly altered mannitol- and NaCl-induced _mt_Ca^2+^ amplitudes, curve area and peak time in *2OX:mtAq* and *mcu*^*1-6*^*:mtAq* plants, our study has identified the critical role for MCUs in maintaining _mt_Ca^2+^ homeostasis *in planta*, and defining the _mt_Ca^2+^ signature. MCU-mediated mitochondrial Ca^2+^ uptake does not appear to substantially affect _cyt_Ca^2+^ dynamics. It is possible that MCU-mediated mitochondrial Ca^2+^ uptake may have a limited capacity to buffer _cyt_Ca^2+^. Alternatively, lack of changes in _cyt_Ca^2+^ may be due to the limitation of our study in which _mt_Ca^2+^ signal responses were recorded at the whole plant level ^40^. Thus, we cannot exclude the possibility that MCUs may buffer _cyt_Ca^2+^ within microdomains. Collectively, our results suggest that MCU family members are functionally redundant in controlling mitochondrial Ca^2+^ uptake. Specific changes in _mt_Ca^2+^ dynamics are caused by enhancing or impairing MCU functions, further highlighting the important roles of MCUs as mitochondrial Ca^2+^ uptake systems in maintaining _mt_Ca^2+^ homeostasis and in sensing changing (micro)environments for mitochondrial function.

The physiological roles of organellar Ca^2+^ transient and homeostasis in organelle functioning are largely unknown ^17^. We show that perturbation of _mt_Ca^2+^ homeostasis apparently interrupts the transcription and translation of mitochondrial genome with impacts on mitochondrial metabolism. We discover that iMUCH activates multi-compartmental UPR and cytosolic translation response, suggesting that iMUCH is likely a pathological condition that evokes the adaptive response as the part of mitochondrial-cytosolic protein quality control mechanisms that have been unified as mitoprotein-induced stress response ^56^. Our study raises questions regarding how perturbation of _mt_Ca^2+^ homeostasis is coordinated with alterations in inter-compartment proteostasis with the physiological consequence, such as, impacts on growth, stress resistance and ageing. Although we are unable to unravel the full complexity of iMUCH evoked adaptive response at this stage, we demonstrate that iMUCH causes misregulation of mtDNA gene expression, and/or disturbance of OXPHOS complex assembly, thus, resulting in a stoichiometric imbalance between nDNA and mtDNA-encoded OXPHOS subunits. A recent study revealed that disturbed chloroplast ion homeostasis impairs plastid rRNA maturation due to the lack of two plastidial envelope-localized K^+^ exchange antiporter (KEA) transporters (KEA1/KEA2) ^57^. Interestingly, loss of KEA1 and KEA2 impaired the osmotic-induced cytosolic Ca^2+^ increase, although it is unknown whether stromal Ca^2+^ dynamics is affected ^58^. Nevertheless, it is an excellent demonstration how the organellar ion homeostasis affects the act of translation machinery within organelles.

The consequence of a mitonuclear protein imbalance has not been fully explored yet in plants, although the study in other systems reveal that mitonuclear protein imbalances triggers a nuclear transcriptional response to ultimately restore mitochondrial proteostasis by activation of the UPR^mt 15^. And in plants interrupting mitochondrial ribosome function was shown to affect mitochondrial gene expression by altering translation and splicing ^60,61^. Here, we show the iMUCH evoked transcriptional response with most upregulated hits of cytosolic ribosome and mitochondrial ribosome, in addition to multi-compartmental chaperones (Fig. 3a-c). Transcriptional activation of cytosolic, mitochondrial, chloroplast and ER chaperone genes highlights the cross-compartmental communication that is likely initiated by iMUCH activated UPR^mt^ that is connected to the UPR response not only in the cytosol (UPR^cyt^), but also in the chloroplast (UPR^chl^) and ER (UPR^er^) (Fig. 8). These results suggest that an intrinsic cellular compartmental communication network operates when mitonuclear protein imbalance occurs, for maintaining cellular homeostasis in plants. This intrinsic communication network is also effective at the post-transcriptional level with a selective translational repression of nDNA *cyRP*s, *mtRP*s, and *cpRP*s, implying that signals derived from iMUCH are not only relayed to the nucleus, but to the cytosol for post-transcriptional regulation of intercompartmental proteostasis (Fig. 8). More than half of all transcriptionally upregulated *cyRP*s are translationally repressed, as evidenced by their reduced translational efficiency and protein abundance. While cyRPs control cytosolic proteostasis, RNA modification proteins likely are key players in tuning organellar proteostasis since these proteins function in organellar RNA editing and translation. Mitoribosomes contain 10 PPR proteins, and rPPR1 is a generic translation factor for mitochondrial translation ^62^. Although rPPR1 was not detected in our proteomic analysis, rPPR9 (At1g60960) was significantly reduced in the sextuple mutant, but not in AA-treated wildtype plants (Supplementary Data 6). These results imply that both RPs and PPR proteins are effectors in iMUCH evoked proteostatic response. Therefore, our findings provide a framework to better understand how a regulatory network that combines transcriptional and post transcriptional mechanisms to coordinate nuclear- and mitochondrial-encoded gene expression for maintaining cellular homeostasis ^63,64^.

**Fig. 8.**
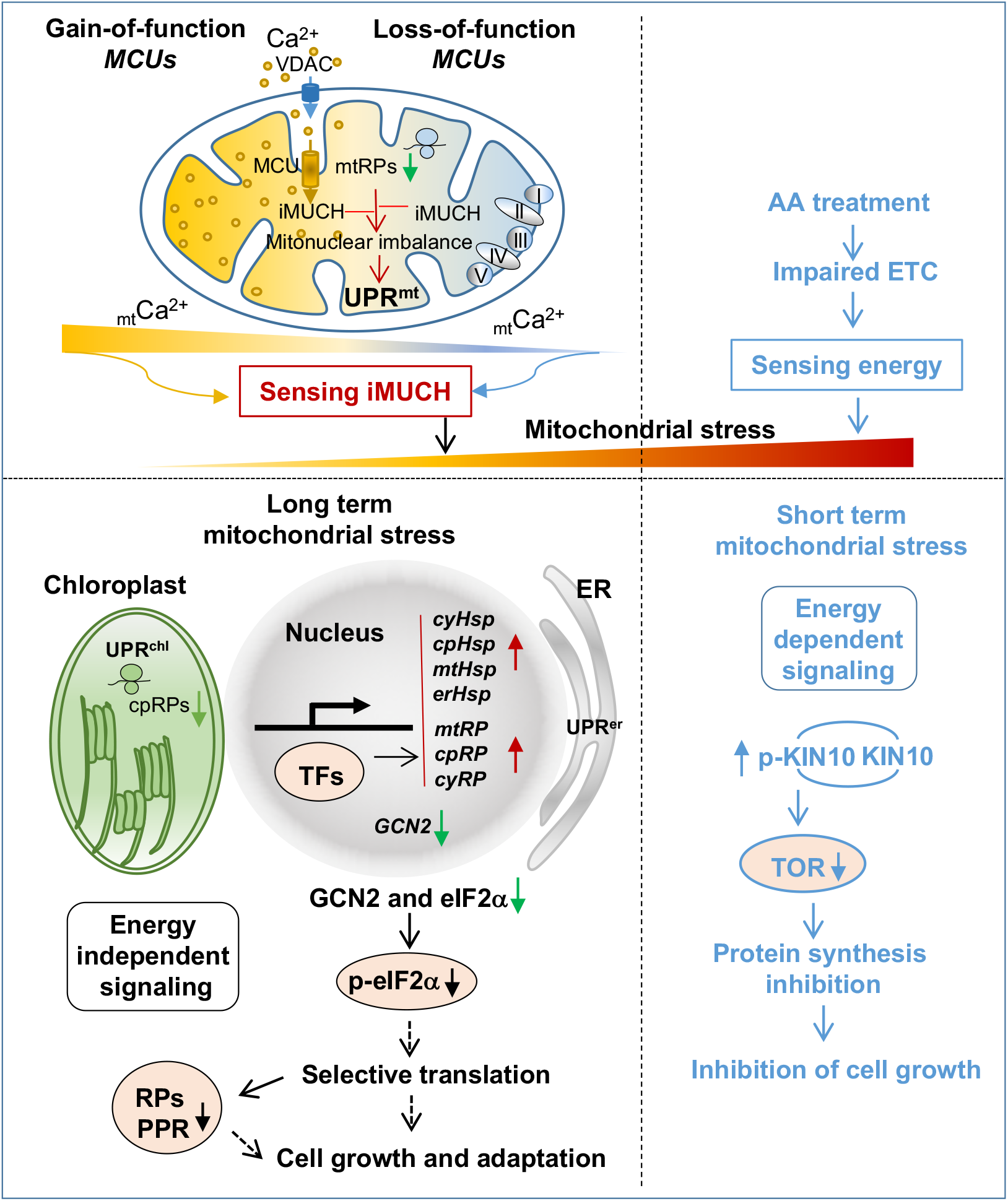
Perturbation of mitochondrial Ca^2+^ homeostasis activates a cross-compartmental proteostatic response. Gain-of-function (*2OX*) or loss-of-function (*sextuple* mutant) MCU proteins impairs mitochondrial Ca^2+^ homeostasis (iMUCH). iMUCH causes misregulation of mtDNA gene expressions that triggers mitonuclear protein imbalance and actives mitochondrial unfolded protein response (UPR^mt^). The signals derived from iMUCH are relayed by unknown transcription factors (TFs) to the nucleus to activate multi-compartmental unfolded protein response (UPR) gene, and ribosome (RP) gene expression. A decrease in global translation with selective translational repression occurs in cytosol upon iMUCH. Reduction of eIF2α phosphorylation (p-eIF2α) is likely attributed to transcriptional and translational inhibition of GCN2 and translational suppression of eIF2α. The selective translational repression of nucleus-encoded mitoribosome and cytoribosome proteins (RPs), and organellar RNA editing pentatricopeptide repeat (PPR) proteins is controlled by an unknown mechanism. Both reduction of eIF2α phosphorylation and selective synthesis of proteins are the part of adaptive mechanisms that protect the cell from severe inhibition of protein synthesis and maintain cellular proteostasis for redirecting cell growth and stress adaptation. AA-impaired respiration electron transfer chain (ETC) causes cellular energy deficiency, thus evokes a short term acute mitochondrial stress that is sensed by KIN10 through phosphorylation of KIN10 (p-KIN10). Increase of p-KIN10 inactivates TOR activity. An energy dependent signalling pathway mediates AA inhibition of cell growth. Together, it is proposed that the GCN2-eIF2α and TOR-S6K pathways mediate mitochondrial stress dependent regulation of protein synthesis by sensing iMUCH and cellular energy status, respectively. Dashed lines indicate the postulated processes, and solid lines indicated defined processes in this study.

Signals and their transducers in iMUCH evoked proteostatic responses are unknown. Our data do not support the notion that reactive oxygen species (ROS) act as the primary signal for the initiation of the transcriptional response because *AOX1a* transcript levels were not significantly raised in *2OX and* sextuple mutant. Likewise, AA treatment induces *AOX1a* expression but cannot elicit the same response. At present, TFs that were upregulated or downregulated upon iMUCH are potential candidates in iMUCH triggered transcriptional pathway (Supplementary Data 1). Phosphorylation of eIF2α has been implicated in translational repression in response to various stresses ^65,66^. We found that *eIF2*α was transcriptionally upregulated, but the level of its encoded protein was significantly reduced. The reduction of p-eIF2α and unchanged TOR level in *2OX* and sextuple mutant is unexpected given that cytosolic translation is to some extent decreased in these plants. A recent study in mammalian cells propose that reduction of eIF2α phosphorylation is an adaptive mechanism that protects the cell from severe inhibition of protein synthesis under long-term mitochondrial stress ^52^. This adaptive mechanism may also apply to the plant with iMUCH, which suffers a long-term mild mitochondrial stress compared to AA induced a short-term acute mitochondrial stress (Fig. 8). An adaptive response to a short-term acute mitochondrial stress via TOR pathway allows the cell to sense cellular energy status to restore proteostasis to avoid cellular death (Fig. 8) ^10,52,59^. Whereas, under a long-term mitochondrial stress, the cell may benefit from eIF2α mediated feedback control of protein synthesis for survival. We assume that eIF2α regulators and targets in plants, like ATF4 and PERK kinase in mammalian cells, are involved in mitochondrial-cytosolic protein quality control mechanisms for coping with mitochondrial stress ^67-70^. Moreover, the selective translation has been considered for aiding recovery from mitochondrial stress ^53,71-74^. Therefore, identifying eIF2α regulators and targets, and putative mechanisms for the selective repression of mRNA translation upon iMUCH will be an exciting area of future work. Nevertheless, we propose that RP and RNA modification proteins identified in this study are effectors in selective translation, which may form a feedback loop to govern capacity of multi-compartment proteostasis. This idea is supported by the following evidence: 1) certain combinations of eIF and RP affect the translation of specific transcripts ^75^; 2) individual ribosome proteins affect translational control by altering ribosome composition ^75^; and 3) PPR, TPR, and PUS proteins are organellar RNA editors and parts of organellar ribosome compositions that are directly involved in organellar translational regulation ^62^. Given the limited genetic accessibility of mitochondria and chloroplasts ^76^, manipulation of ribosome and ETC complexes encoded by the mitochondrion and chloroplast genomes is feasible, but still very changeling ^77-80^. Our study demonstrates that mtDNA *mtRPs* and cpDNA *cpRP*s are selectively repressed upon iMUCH, thus providing an excellent system for studying the regulation of mitochondrial and chloroplast gene transcription and translation.

Our findings in plants are merged with increasing evidence from studies in yeast and animals for supporting that mitochondrial stress dependent regulation of protein synthesis has impacts on growth, stress resistance and ageing ^56^. iMUCH is linked to the inhibition of cell growth, which is a phenotype reminiscent of the inhibition of cell proliferation ^81^. The effects of iMUCH on growth are different from AA mediated growth inhibition that is controlled by TOR pathway in sensing cellular energy status (Fig. 8). Adaptive actions of the cell upon iMUCH may also include transcriptional and post-transcriptional co-suppression of cell growth pathways that involve growth hormones and cell wall biogenesis. On the other hand, iMUCH increases the vulnerability to cellular senescence and stress. Cellular senescence is triggered by numerous stressors and physiological processes, including mitochondrial dysfunction, perturbed proteostasis and ribosomal stress etc. ^82^. One of the hallmarks of the senescence is deregulated metabolism, which is also evident in 2OX and sextuple mutant. Whether other inter-dependent features, for example, cell-cycle withdraw, macromolecular damage and secretory phenotype mark 2OX and sextuple mutant is an interesting question for the future study. Nevertheless, iMUCH offers a pathological condition for studying how mitochondria regulate cellular senescence *in vivo* ^82^. While 2OX and sextuple mutant show similar phenotypes in growth and stress resistance, the senescence occurs in relative late life stage in sextuple mutant compared to 2OX. Given that the timing of senescence in 2OX plants is related to *MCU2* expression level, it is assumed that increasing mitochondrial Ca^2+^ uptake may accelerate cell-cycle arrest that is initiated by iMUCH. Therefore, 2OX plants show much earlier and severe senescence than sextuple mutant.

In summary, our study provides molecular and genetic evidence for the essential functions of MCU proteins in maintaining _mt_Ca^2+^ homeostasis to protect mitochondria from stresses that normally trigger mitochondrial Ca^2+^ uptake. Understanding iMUCH induced adaptive mechanism governing growth, stress response and senescence should bring great promise to maximize plant production under stress conditions.

## Methods

### Plant materials and general growth conditions

All wildtype, mutant, and transgenic Arabidopsis (*Arabidopsis thaliana*) lines used in this study were in the Columbia (Col-0) accession. Arabidopsis *MCU1* (At1g09575), *MCU2* (At1g57610), *MCU3* (AT2G23790), *MCU4* (At4g36820), *MCU5* (At5g42610), and *MCU6* (At5g66650) cDNAs were amplified by RT-PCR and cloned in-frame upstream of *YFP* into the binary vector pPZP200. About 1000bp fragments encompassing sequence upstream of the ATG were amplified from each *MCU* gene, and cloned into binary vector pMDC162. Transgenic plants expressing *35S:MCU1-YFP* and *proMCU1:GUS, 35S:MCU2-YFP* and *proMCU2:GUS, 35S:MCU3-YFP* and *proMCU3:GUS, 35S:MCU4-YFP* and *proMCU4:GUS, 35S:MCU5-YFP* and selected on medium containing 50 μg/ml gentamycin (*35S:MCU:YFP*) and 20 μg/ml hygromycin (*proMCU:GUS)* containing plates. Single mutants *mcu1-1* (SALK_082151), *mcu2-1* (SALK_011710), *mcu3-1* (SALK_019312), *mcu4-1* (SALK_036975), and *mcu6-1* (SALK_037347) were obtained from the Arabidopsis Biological Resource Center (ABRC), and *mcu5-1* was generated using CRISPR/Cas9-mediated genome editing as previously described (Liu et al., 2015). The higher-order mutant *mcu1-1 mcu2-1 mcu3-1 mcu4-1 mcu5-1 mcu6-1* was generated using genetic crosses followed by identification of homozygous mutant plants from F_2_ or F_3_ progeny using PCR-based genotyping. Six MCU single mutants in a *mtAeq* line were generated by CRISPR/Cas9 editing approach using two single guide RNAs (sgRNA1 and sgRNA2) targeting sites within *MCU1, MCU2, MCU3, MCU4, MCU5*, and *MCU6* to obtain the homozygous *mcu1aq, mcu2aq, mcu3aq, mcu4aq, mcu5aq*, and *mcu6aq* single mutants with a single copy of deletion, respectively.

### Subcellular localization of MCU proteins and GUS staining

Subcellular localization of MCU proteins was examined using confocal microscopy (Zeiss 710) in the leaf and root cells of stable *35S:MCU-YFP* transgenic plants. YFP and chlorophyll fluorescence were excited at 514 nm with detection ranges of 525–600 nm and 650–720 nm for YFP and chlorophyll fluorescence, respectively. MitoTracker dye was excited at 644 nm with a detection range of 578-696 nm for MitoTracker™ Deep Red FM (Invitrogen M22426). For GUS staining, seedlings expressing *proMCUs:GUS* were submerged in the staining solution and incubated at room temp or 37°C until blue developed.

### RNA isolation for cDNA amplification and RT-qPCR

Total RNA was extracted from less than 100 mg of 10-day-old seedlings that were grown vertically on MS plates, or treated with 50 μM AA or DMSO for 4 hours in liquid MS. First-strand cDNA was synthesized using the HiScript^®^ II cDNA Synthesis Kit (Vazyme R212-02), and oligo d(T) for the synthesis of cDNA used for amplification of nDNA encoded genes, and random primers for amplification of mtDNA encoded genes, respectively. qPCR with iTaq Universal SYBR^®^ Green Supermix (BIORAD) was conducted on a Roche 480 Real-Time PCR system following the manufacturer’s instructions.

### Aequorin-based FAS imaging of _mt_Ca^2+^ and _cyt_Ca^2+^ response

For luminescence imaging of _mt_Ca^2+^ and _cyt_Ca^2+^, the two constructs *Ubpro:CytoAequorin-YFP* (*cytAeq*, CD3-1798) and *Ubpro:MitoAequorin-YFP* (*mtAeq*, CD3-1800) were used, consisting of the *Ubiquitin* promoter driving the expression of aequorin targeted to the cytosol and mitochondrial matrix, respectively ^40,83^(Mehlmer et al., 2012; Zhu *et al*., 2013). These constructs were transformed into wildtype Col-0, *MCU2* overexpressing (*2OX*), and *mcu1-1 mcu2-1 mcu3-1 mcu4-1 mcu5-1 mcu6-1* sextuple mutant plants. Homozygous plants harboring *cytAeq* and *mtAeq* were subjected to salt and mannitol stimuli to record _mt_Ca^2+^ and _cyt_Ca^2+^ response using the aequorin-based FAS recording system ^40^(Zhu *et al*., 2013). Briefly, 8-day-old seedlings grown vertically on MS plates were transferred onto an adhesive film. Film-adhesive seedlings (FAS) were incubated with coelenterazine solution (final concentration 5 μmol/L) in the dark for at least 5 hours at room temperature. The Ca^2+^ luminescence image with a 60s integration of emitted light was then acquired immediately after FAS was subjected to 100mM NaCl or 400mM mannitol stimuli using a CCD camera (NightShade LB 985). The discharge luminescence image with a 60s integration of emitted light was acquired immediately after FAS was subjected to discharge buffer (20% (v/v) ethanol solution containing 1M Ca_2_Cl). Luminescence intensity of each FAS film containing about 30 seedlings was quantified by indiGO™ software, and converted to Ca^2+^ concentration by the formula pCa=0.332588(-logk)+5.5593, where k is a rate constant equal to stimulus-induced Ca^2+^ luminescence counts (CaL) divided by discharge luminescence counts (DisL), k=(CaL)/(DisL+CaL) ^40,84^. For Ca^2+^ luminescence imaging by FAS assay, more than 6 independently prepared films for each genotype were recorded.

### Luminometry reading of _mt_Ca^2+^ and _cyt_Ca^2+^ response

For quantitative measurement of Ca^2+^ response using a luminometer (GLOMAX, Promega, USA, Model: 2031-002, Serial: 203001553), the 1.5 mL tube containing a single whole 8-day-old seedling incubated with coelenterazine solution (final concentration 5 μmol/L) for overnight was set on the chamber for recording of the resting luminescence for about 60 s. Ca^2+^ luminescence value was read at 1 s interval for another 100 s starting by adding a final concentration of 100 mM NaCl or 400 mM mannitol into the tube. The remaining aequorin was discharged by adding discharge buffer, and luminescence value was read for 100 s. The _mt_Ca^2+^ dynamics of the *mcu1aq, mcu2aq, mcu3aq, mcu4aq, mcu5aq*, and *mcu6aq* single mutants, wildtype, 2OX and sextuple mutant seedlings were performed using a luminometer. The luminescence value of each individual seedling was converted to Ca^2+^ concentration by the formula described above. For Ca^2+^ luminescence reading by luminometer, 10–15 whole seedlings were independently measured. Ca^2+^ amplitudes, curve area, peak time, were calculated by comparing the mannitol or NaCl-induced _mt_[Ca^2+^]^peak^ against that of _mt_[Ca^2+^]^rest^. Average _mt_Ca^2+^ amplitude, curve area and peak time were calculated from the data of independent measurement of 10–15 seedlings for each genotype. The area of _mt_Ca^2+^ response curve was determined by Graphpad Prism 9.

### RNA-seq and data analysis

Wildtype (Col-0), *2OX, 4OX, 6OX*, and sextuple mutant seedlings were grown vertically on half-strength MS plates. Ten-day-old whole seedlings were then transferred into 9-well plates containing half-strength liquid MS medium overnight with shaking in the dark, then treated with 50 μM antimycin A (AA) or DMSO for 4 h. Total RNA from AA-treated and DMSO-treated control seedlings was prepared using Trizol (Invitrogen). Sequencing libraries were generated from 1 μg RNA per sample using NEBNext® Ultra™ RNA Library Prep Kit for Illumina® (NEB, USA) following the manufacturer’s recommendations. The reads were then mapped against the *Arabidopsis thaliana* reference genome (TAIR10). Cufflinks v2.2.1 was used to obtain normalized reads per kilobase of transcript per million mapped reads (RPKM) values for each annotated gene. Three biological replicates of each treatment were used for RNA-seq analysis.

### GO enrichment analysis

The PANTHER website was used for GO term enrichment analysis of upregulated or downregulated genes (fold-change > 2, Benjamini-Hochberg adjusted *p*-value < 0.01) in RNA-seq data from Col-0, *2OX, 4OX, 6OX*, and the sextuple mutant treated with 50 μM antimycin A. imageGP was used to generate GO enrichment plots and heatmaps. RNA-seq volcano plots were created in R using the *ggplot2* package (Wickham, 2016)

### Immunoblotting

For preparation of total proteins, 10-day-old seedlings that were grown vertically on MS plates, or treated with 50 μM AA or DMSO in liquid MS for 4 h (∼200 mg) were collected and ground to a fine powder in liquid nitrogen, then homogenized with two volumes of extraction buffer containing 125 mM Tris-HCl, pH 8.8, 1% (w/v) SDS, 10% (v/v) glycerol, and 50 mM Na_2_S_2_O_2_, 0.2 mM PMSF and protease inhibitor cocktail (Roche). The homogenates were incubated on ice for 5 min and centrifuged at 13,000*g* for 10 min at 4ºC. The supernatants were separated by 12.5% SDS-PAGE, transferred to Immobilon-P^SQ^ positively charged nylon membrane, and immunoblotted with antibodies purchased from Agrisera and PhytoAB (Supplementary Data 10).

For preparation of enriched mitochondrial proteins, 10-day-old seedlings (∼5 g) were ground in liquid nitrogen and extracted in 30 mL extraction buffer (15 mM MOPs, pH 7.4, 450 mM sucrose, 0.6% [w/v] PVP40, 0.2% [w/v] BSA, 1.5 mM EGTA, 10 mM DTT, 0.2 mM PMSF and protease inhibitor cocktail). Homogenates were then filtered through two layers of Miracloth and centrifuged at 3,500*g* for 10 min, followed by additional centrifugations at 8,000*g* for 5 min and at 17,000*g* for 30 min at 4ºC. The resulting pellets were washed with 1 mL washing buffer (0.3 M sucrose, 15 mM MOPS, pH 7.2) twice. Pellets were resuspended in 200 μL extraction buffer for immunoblotting with antibodies purchased from Agrisera and PhytoAB (Supplementary Data 10).

### Proteomic analysis

Ten-day-old whole seedlings of wildtype (Col-0) and sextuple mutant seedlings grown vertically on half-strength MS plates were collected. For AA treatment, ten-day-old whole seedlings of Col-0 were then transferred into 9-well plates containing half-strength liquid MS medium overnight with shaking in the dark, then treated with 50 μM antimycin A (AA) or DMSO for 4 h. Whole seedlings (100mg) were homogenized in lysis buffer consisted of 2.5% SDS/100 mM Tris-HCl (pH 8.0). Then the samples were subjected to ultra-sonication. After centrifugation, proteins in the supernatant were precipitated by adding 4 times of pre-cooled acetone. The protein pellet was dissolved in 8 M Urea/100 mM Tris-Cl. After centrifugation, the supernatant was used for reduction reaction (10 mM DTT, 37ºC for 1 h), and followed by alkylation reaction (40 mM iodoacetamide, room temperature/dark place for 30 min). Urea was diluted below 2 M using 100 mM Tris-HCl (pH 8.0). Trypsin was added at a ratio of 1:50 (enzyme:protein, w/w) for overnight digestion at 37ºC. TMT labeling was performed according to manufacturer’s instructions. LC-MS/MS data acquisition was carried out on an Orbitrap Exploris 480 mass spectrometer coupled with an Easy-nLC 1200 system. MS raw data were analyzed with MaxQuant (V1.6.6) using the Andromeda database search algorithm. Spectra files were searched against the UniProt Arabidopsis thaliana proteome database. Search results were filtered with 1% FDR at both protein and peptide levels.

### Polysome profiling and TE calculations

Sample preparation, collection and AA treatment are the same as described in proteomic analysis. Whole seedlings (∼200 mg) were ground in liquid nitrogen and extracted in 1 mL polysome buffer (400 mM Tris-HCl, pH 8.4, 100 mM MgCl_2_, 200 mM KCl, 0.5% [v/v] NP-40, 50 μM EGTA, 50 μg/mL cycloheximide, 50 μg/mL chloramphenicol, and 400 U/mL recombinant RNasin [RNase inhibitor, Promega]). The samples were centrifuged at 12,000 g for 10 min at 4ºC to remove debris. The supernatant for each sample (1 mL) was loaded on top of the sucrose gradient. Gradients of 20–50% sucrose (11 mL) in gradient buffer (40 mM Tris-HCl, pH 8.4, 10 mM MgCl_2_, 20 mM KCl) were prepared in polypropylene (14 × 89 mm) centrifuge tubes (Seton, CAN). The gradients were ultra-centrifuged for 3 h at 175,000*g* in a SW41Ti rotor (Beckman-Coulter, USA) and analyzed by measuring continuous absorbance at 260 nm using a Piston Gradient Fractionator (Biocomp Instruments). Total RNA of polysome or monosome fractions was precipitated and pooled for RNA-seq analysis ^85,86^(Missra and von Arnim, 2014; Mustroph et al., 2009). Sequencing libraries were generated using NEBNext® Ultra™ RNA Library Prep Kit for Illumina® (NEB, USA) as described above. Data analysis was performed as described previously ^50^(Molenaars *et al*., 2020).

TEs were calculated as the ratio between the transcripts in the polysome fractions (highly translated ribosome-mRNA fractions) and the transcripts in the monosome fractions (lower translated ribosome-mRNA fractions) ^50^(Molenaars *et al*., 2020). TEs were then compared between *2OX* or sextuple mutant samples and wildtype control samples as defined: log_2_ [(MCU^polysome^/MCU^monosome^) / (WT^polysome^/WT^monosome^)]. Monosome and polysome fractions were collected as indicated in Fig. 4. Means of three replicates were used to calculate the ratios.

### Metabolic analysis of energy-related metabolites

Sample preparation, collection and AA treatment are the same as described in proteomic analysis. Whole seedlings (50mg) were accurately weighed and placed in a 2 mL EP tube, then added 500 μL pre-cooled extractant (70% methanol aqueous solution). Homogenize 4 times at 30 Hz for 30 s each time. Shake at 1500 r / min for 5 min, then stand for 15 min on ice; and centrifuge with 12000 r/min at 4ºC for 10 min. Finally take 200 μL of supernatant into the sample bottle for LC-MS/MS analysis. The extracts were analyzed using an LC-ESI-MS/MS system (UPLC, Shim-pack UFLC SHI-MADZU CBM30A system, https://www.shimadzu.com/ MS, QTRAP® System, https://sciex.com/). LIT and triple quadrupole (QQQ) scans were acquired on a triple quadrupole-linear ion trap mass spectrometer (QTRAP), QTRAP® LC-MS/MS System, equipped with an ESI Turbo Ion-Spray interface, operating in positive and negative ion mode and controlled by Analyst 1.6.3 software (Sciex). Unsupervised PCA (principal component analysis) was performed by statistics function prcomp within R (www.r-project.org). Significantly regulated metabolites between groups were determined by VIP >= 1 and absolute Log2FC (fold change) >= 1. VIP values were extracted from OPLS-DA result, which also contain score plots and permutation plots, was generated using R package MetaboAnalystR.

### Growth assessment under stress

Arabidopsis seeds were surface-sterilized and sown on solidified half Murashige and Skoog (MS) medium plates. After stratification at 4ºC for 2 d in the dark, the plates were placed vertically at 23ºC, under long-day (16-h light and 8-h dark) photoperiodic condition. Five-day-old seedlings were transferred to MS plates containing 150 mM mannitol. In addition, 10-day-old seedlings were transferred to soil for subsequent growth at 23ºC in a growth room under long-day photoperiodic conditions as described above.

### Data availability

The RNA-seq and Ribo-seq data generated in this study have been deposited to Gene Expression Omnibus (GEO) under the accession number GSE186418. The mass spectrometry proteomics data have been deposited to the ProteomeXchange Consortium (http://proteomecentral.proteomexchange.org) via the iProX partner repository ^87^ with the Table identifier PXD029171.The energy metabolite data were deposited to MetaboLights ^87,88^ under the Table identifier MTBLS3621 (www.ebi.ac.uk/metabolights/MTBLS3621).

## Acknowledgements

This work was supported by a special support program from Henan University and by a grant from the National Natural Science Foundation of China to X.Z. (31872807), and by the Chinese Academy of Sciences, the CAS Strategic Priority Research Program to J.-K.Z. (XDB27040101). We thank Dr. Pengcheng Wang and Dr. Hairong Zhang for providing us antibodies.

## Author contributions

X.Z. and J.-K.Z. conceived the project and wrote the article. X.Z., C.M. and X.B. generated transgenic plants, performed Ca^2+^ imaging and completed molecular biology and biochemistry experiments. S.Z., Z.O. and Z.Z. participated in generating transgenic plants. K.T. and S.X. conducted RNA-seq analysis. Y.W., D.Z, H.X., Q. L, X.W. and J. Z. assisted with experiments. X.Z., J.-K.Z. C.-P.S. and S.P. D.-K analyzed the data.

## Supplemental data

**Supplementary Data 1. Differentially expressed genes (DEGs) identified in 2OX and sextuple mutant.**

**Supplementary Data 2. A list of upregulated DEGs enriched in ribosome and translation**.

**Supplementary Data 3. Differentially expressed chaperone genes**.

**Supplementary Data 4. Differentially expressed genes (DEGs) identified in AA-treated 2OX and sextuple mutant**.

**Supplementary Data 5. A list of AA activated transcription factors (GO 0140110).**

**Supplementary Data 6. Mass spectrometry proteomics data**.

**Supplementary Data 7. The list of genes for proteome–transcriptome cross-analysis.**

**Supplementary Data 8. RNA-seq data of the monosomal and polysomal RNA for analysis of translation efficiency**.

**Supplementary Data 9. Metabolic profiling of energy metabolites.**

**Supplementary Data 10. Primer sets and antibodies**.

**Supplementary Data 11. Statistics information**.

## References

1 Zhu, J. K. Abiotic Stress Signaling and Responses in Plants. Cell 167, 313–324, doi:10.1016/j.cell.2016.08.029 (2016).

2 Van Aken, O. et al. Defining the mitochondrial stress response in Arabidopsis thaliana. Molecular plant 2, 1310–1324, doi:10.1093/mp/ssp053 (2009).

3 Butow, R. A. & Avadhani, N. G. Mitochondrial signaling: The retrograde response. Molecular cell 14, 1–15, doi:Doi 10.1016/S1097-2765(04)00179-0 (2004).

4 de Souza, A., Wang, J. Z. & Dehesh, K. Retrograde Signals: Integrators of Interorganellar Communication and Orchestrators of Plant Development. Annual Review of Plant Biology, Vol 68 68, 85–108, doi:10.1146/annurev-arplant-042916-041007 (2017).

5 Quiros, P. M., Mottis, A. & Auwerx, J. Mitonuclear communication in homeostasis and stress. Nature reviews. Molecular cell biology 17, 213–226, doi:10.1038/nrm.2016.23 (2016).

6 Mottis, A., Herzig, S. & Auwerx, J. Mitocellular communication: Shaping health and disease. Science 366, 827–832, doi:10.1126/science.aax3768 (2019).

7 Andreasson, C., Ott, M. & Buttner, S. Mitochondria orchestrate proteostatic and metabolic stress responses. EMBO reports 20, e47865, doi:10.15252/embr.201947865 (2019).

8 Kim, H. E. et al. Lipid Biosynthesis Coordinates a Mitochondrial-to-Cytosolic Stress Response. Cell 166, 1539–1552 e1516, doi:10.1016/j.cell.2016.08.027 (2016).

9 D’Amico, D., Sorrentino, V. & Auwerx, J. Cytosolic Proteostasis Networks of the Mitochondrial Stress Response. Trends in biochemical sciences 42, 712–725, doi:10.1016/j.tibs.2017.05.002 (2017).

10 Wang, X. & Chen, X. J. A cytosolic network suppressing mitochondria-mediated proteostatic stress and cell death. Nature 524, 481–484, doi:10.1038/nature14859 (2015).

11 Wrobel, L. et al. Mistargeted mitochondrial proteins activate a proteostatic response in the cytosol. Nature 524, 485–488, doi:10.1038/nature14951 (2015).

12 Boos, F. et al. Mitochondrial protein-induced stress triggers a global adaptive transcriptional programme. Nat Cell Biol 21, 442–451, doi:10.1038/s41556-019-0294-5 (2019).

13 Hartl, F. U., Bracher, A. & Hayer-Hartl, M. Molecular chaperones in protein folding and proteostasis. Nature 475, 324–332, doi:10.1038/nature10317 (2011).

14 Horwich, A. L. Molecular chaperones in cellular protein folding: the birth of a field. Cell 157, 285–288, doi:10.1016/j.cell.2014.03.029 (2014).

15 Houtkooper, R. H. et al. Mitonuclear protein imbalance as a conserved longevity mechanism. Nature 497, 451–457, doi:10.1038/nature12188 (2013).

16 Logan, D. C. & Knight, M. R. Mitochondrial and cytosolic calcium dynamics are differentially regulated in plants. Plant physiology 133, 21–24, doi:10.1104/pp.103.026047 (2003).

17 Resentini, F., Ruberti, C., Grenzi, M., Bonza, M. C. & Costa, A. The signatures of organellar calcium. Plant physiology 187, 1985–2004, doi:10.1093/plphys/kiab189 (2021).

18 Griffiths, E. J. & Rutter, G. A. Mitochondrial calcium as a key regulator of mitochondrial ATP production in mammalian cells. Biochimica et biophysica acta 1787, 1324–1333, doi:10.1016/j.bbabio.2009.01.019 (2009).

19 Giacomello, M., Drago, I., Pizzo, P. & Pozzan, T. Mitochondrial Ca2+ as a key regulator of cell life and death. Cell death and differentiation 14, 1267–1274, doi:10.1038/sj.cdd.4402147 (2007).

20 Calvo-Rodriguez, M. et al. Increased mitochondrial calcium levels associated with neuronal death in a mouse model of Alzheimer’s disease. Nature communications 11, 2146, doi:10.1038/s41467-020-16074-2 (2020).

21 Bernardi, P. & Rasola, A. Calcium and cell death: the mitochondrial connection. Sub-cellular biochemistry 45, 481–506, doi:10.1007/978-1-4020-6191-2_18 (2007).

22 Garbincius, J. F. & Elrod, J. W. Mitochondrial Calcium Exchange in Physiology and Disease. Physiol Rev 102, 893–992, doi:10.1152/physrev.00041.2020 (2022).

23 Giorgio, V., Guo, L., Bassot, C., Petronilli, V. & Bernardi, P. Calcium and regulation of the mitochondrial permeability transition. Cell Calcium 70, 56–63, doi:10.1016/j.ceca.2017.05.004 (2018).

24 Rizzuto, R., De Stefani, D., Raffaello, A. & Mammucari, C. Mitochondria as sensors and regulators of calcium signalling. Nature reviews. Molecular cell biology 13, 566–578, doi:10.1038/nrm3412 (2012).

25 Baughman, J. M. et al. Integrative genomics identifies MCU as an essential component of the mitochondrial calcium uniporter. Nature 476, 341–345, doi:10.1038/nature10234 (2011).

26 Liu, J. C. et al. EMRE is essential for mitochondrial calcium uniporter activity in a mouse model. JCI insight 5, doi:10.1172/jci.insight.134063 (2020).

27 De Stefani, D., Raffaello, A., Teardo, E., Szabo, I. & Rizzuto, R. A forty-kilodalton protein of the inner membrane is the mitochondrial calcium uniporter. Nature 476, 336–340, doi:10.1038/nature10230 (2011).

28 Sancak, Y. et al. EMRE is an essential component of the mitochondrial calcium uniporter complex. Science 342, 1379–1382, doi:10.1126/science.1242993 (2013).

29 Hoffman, N. E. et al. MICU1 motifs define mitochondrial calcium uniporter binding and activity. Cell reports 5, 1576–1588, doi:10.1016/j.celrep.2013.11.026 (2013).

30 Kamer, K. J. & Mootha, V. K. MICU1 and MICU2 play nonredundant roles in the regulation of the mitochondrial calcium uniporter. EMBO reports 15, 299–307, doi:10.1002/embr.201337946 (2014).

31 Wang, Y. et al. Structural Mechanism of EMRE-Dependent Gating of the Human Mitochondrial Calcium Uniporter. Cell 177, 1252–1261 e1213, doi:10.1016/j.cell.2019.03.050 (2019).

32 Fan, M. et al. Structure and mechanism of the mitochondrial Ca(2+) uniporter holocomplex. Nature 582, 129–133, doi:10.1038/s41586-020-2309-6 (2020).

33 Wagner, S. et al. The EF-Hand Ca2+ Binding Protein MICU Choreographs Mitochondrial Ca2+ Dynamics in Arabidopsis. The Plant cell 27, 3190–3212, doi:10.1105/tpc.15.00509 (2015).

34 Teardo, E. et al. Physiological Characterization of a Plant Mitochondrial Calcium Uniporter in Vitro and in Vivo. Plant physiology 173, 1355–1370, doi:10.1104/pp.16.01359 (2017).

35 Selles, B., Michaud, C., Xiong, T. C., Leblanc, O. & Ingouff, M. Arabidopsis pollen tube germination and growth depend on the mitochondrial calcium uniporter complex. The New phytologist 219, 58–65, doi:10.1111/nph.15189 (2018).

36 Teardo, E. et al. A chloroplast-localized mitochondrial calcium uniporter transduces osmotic stress in Arabidopsis. Nature plants 5, 581–588, doi:10.1038/s41477-019-0434-8 (2019).

37 Carraretto, L. et al. Ion Channels in Plant Bioenergetic Organelles, Chloroplasts and Mitochondria: From Molecular Identification to Function. Molecular plant 9, 371–395, doi:10.1016/j.molp.2015.12.004 (2016).

38 Ruberti, C. et al. MCU proteins dominate in vivo mitochondrial Ca2+ uptake in Arabidopsis roots. The Plant cell, doi:10.1093/plcell/koac242 (2022).

39 Knight, H., Trewavas, A. J. & Knight, M. R. Calcium signalling in Arabidopsis thaliana responding to drought and salinity. The Plant journal : for cell and molecular biology 12, 1067–1078, doi:10.1046/j.1365-313x.1997.12051067.x (1997).

40 Zhu, X., Feng, Y., Liang, G., Liu, N. & Zhu, J. K. Aequorin-based luminescence imaging reveals stimulus- and tissue-specific Ca2+ dynamics in Arabidopsis plants. Molecular plant 6, 444–455, doi:10.1093/mp/sst013 (2013).

41 McAinsh, M. R. & Pittman, J. K. Shaping the calcium signature. New Phytologist 181, 275–294, doi:10.1111/j.1469-8137.2008.02682.x (2009).

42 Loro, G. et al. Targeting of Cameleons to various subcellular compartments reveals a strict cytoplasmic/mitochondrial Ca(2)(+) handling relationship in plant cells. The Plant journal : for cell and molecular biology 71, 1–13, doi:10.1111/j.1365-313X.2012.04968.x (2012).

43 Huang, L. S., Cobessi, D., Tung, E. Y. & Berry, E. A. Binding of the respiratory chain inhibitor antimycin to the mitochondrial bc(1) complex: A new crystal structure reveals an altered intramolecular hydrogen-bonding pattern. J Mol Biol 351, 573–597, doi:10.1016/j.jmb.2005.05.053 (2005).

44 Wikstrom, M. K. & Berden, J. A. Oxidoreduction of cytochrome b in the presence of antimycin. Biochimica et biophysica acta 283, 403–420, doi:10.1016/0005-2728(72)90258-7 (1972).

45 Slater, E. C. The mechanism of action of the respiratory inhibitor, antimycin. Biochimica et biophysica acta 301, 129–154, doi:10.1016/0304-4173(73)90002-5 (1973).

46 Ng, S. et al. A membrane-bound NAC transcription factor, ANAC017, mediates mitochondrial retrograde signaling in Arabidopsis. The Plant cell 25, 3450–3471, doi:10.1105/tpc.113.113985 (2013).

47 Ivanova, A. et al. A Functional Antagonistic Relationship between Auxin and Mitochondrial Retrograde Signaling Regulates Alternative Oxidase1a Expression in Arabidopsis. Plant physiology 165, 1233–1254, doi:10.1104/pp.114.237495 (2014).

48 De Clercq, I. et al. The membrane-bound NAC transcription factor ANAC013 functions in mitochondrial retrograde regulation of the oxidative stress response in Arabidopsis. The Plant cell 25, 3472–3490, doi:10.1105/tpc.113.117168 (2013).

49 Van Aken, O., Zhang, B., Law, S., Narsai, R. & Whelan, J. AtWRKY40 and AtWRKY63 modulate the expression of stress-responsive nuclear genes encoding mitochondrial and chloroplast proteins. Plant physiology 162, 254–271, doi:10.1104/pp.113.215996 (2013).

50 Molenaars, M. et al. A Conserved Mito-Cytosolic Translational Balance Links Two Longevity Pathways. Cell metabolism 31, 549-+, doi:10.1016/j.cmet.2020.01.011 (2020).

51 Antonicka, H. et al. A High-Density Human Mitochondrial Proximity Interaction Network. Cell metabolism 32, 479-+, doi:10.1016/j.cmet.2020.07.017 (2020).

52 Samluk, L. et al. Cytosolic translational responses differ under conditions of severe short-term and long-term mitochondrial stress. Mol Biol Cell 30, 1864–1877, doi:10.1091/mbc.E18-10-0628 (2019).

53 Topf, U., Uszczynska-Ratajczak, B. & Chacinska, A. Mitochondrial stress-dependent regulation of cellular protein synthesis. Journal of cell science 132, doi:ARTN jcs226258 10.1242/jcs.226258 (2019).

54 Merkwirth, C. et al. Two Conserved Histone Demethylases Regulate Mitochondrial Stress-Induced Longevity. Cell 165, 1209–1223, doi:10.1016/j.cell.2016.04.012 (2016).

55 Wang, X. & Auwerx, J. Systems Phytohormone Responses to Mitochondrial Proteotoxic Stress. Molecular cell 68, 540–551 e545, doi:10.1016/j.molcel.2017.10.006 (2017).

56 Boos, F., Labbadia, J. & Herrmann, J. M. How the Mitoprotein-Induced Stress Response Safeguards the Cytosol: A Unified View. Trends in cell biology 30, 241–254, doi:10.1016/j.tcb.2019.12.003 (2020).

57 DeTar, R. A. et al. Loss of inner-envelope K+/H+ exchangers impairs plastid rRNA maturation and gene expression (vol 33, pg 2479, 2021). The Plant cell 33, 3746–3746, doi:10.1093/plcell/koab209 (2021).

58 Stephan, A. B., Kunz, H. H., Yang, E. & Schroeder, J. I. Rapid hyperosmotic-induced Ca2+ responses in Arabidopsis thaliana exhibit sensory potentiation and involvement of plastidial KEA transporters. Proceedings of the National Academy of Sciences of the United States of America 113, E5242–E5249, doi:10.1073/pnas.1519555113 (2016).

59 Topf, U. et al. Quantitative proteomics identifies redox switches for global translation modulation by mitochondrially produced reactive oxygen species. Nature communications 9, doi:ARTN 324 10.1038/s41467-017-02694-8 (2018).

60 Kwasniak, M. et al. Silencing of the Nuclear RPS10 Gene Encoding Mitochondrial Ribosomal Protein Alters Translation in Arabidopsis Mitochondria. The Plant cell 25, 1855–1867, doi:10.1105/tpc.113.111294 (2013).

61 Kwasniak-Owczarek, M., Kazmierczak, U., Tomal, A., Mackiewicz, P. & Janska, H. Deficiency of mitoribosomal S10 protein affects translation and splicing in Arabidopsis mitochondria. Nucleic Acids Res 47, 11790–11806, doi:10.1093/nar/gkz1069 (2019).

62 Waltz, F. et al. Small is big in Arabidopsis mitochondrial ribosome. Nature plants 5, 106–117, doi:10.1038/s41477-018-0339-y (2019).

63 Couvillion, M. T., Soto, I. C., Shipkovenska, G. & Churchman, L. S. Synchronized mitochondrial and cytosolic translation programs. Nature 533, 499-+, doi:10.1038/nature18015 (2016).

64 St-Pierre, J. & Topisirovic, I. Nucleus to Mitochondria: Lost in Transcription, Found in Translation. Developmental cell 37, 490–492, doi:10.1016/j.devcel.2016.06.003 (2016).

65 Wang, L. J. et al. The inhibition of protein translation mediated by AtGCN1 is essential for cold tolerance in Arabidopsis thaliana. Plant Cell Environ 40, 56–68, doi:10.1111/pce.12826 (2017).

66 Lokdarshi, A. et al. Light Activates the Translational Regulatory Kinase GCN2 via Reactive Oxygen Species Emanating from the Chloroplast. The Plant cell 32, 1161–1178, doi:10.1105/tpc.19.00751 (2020).

67 Harding, H. P. et al. Regulated translation initiation controls stress-induced gene expression in mammalian cells. Molecular cell 6, 1099–1108, doi:Doi 10.1016/S1097-2765(00)00108-8 (2000).

68 Quiros, P. M. et al. Multi-omics analysis identifies ATF4 as a key regulator of the mitochondrial stress response in mammals. J Cell Biol 216, 2027–2045, doi:10.1083/jcb.201702058 (2017).

69 Khan, N. A. et al. mTORC1 Regulates Mitochondrial Integrated Stress Response and Mitochondrial Myopathy Progression. Cell Metab 26, 419–428 e415, doi:10.1016/j.cmet.2017.07.007 (2017).

70 Kuhl, I. et al. Transcriptomic and proteomic landscape of mitochondrial dysfunction reveals secondary coenzyme Q deficiency in mammals. Elife 6, doi:10.7554/eLife.30952 (2017).

71 Shi, Z. et al. Heterogeneous Ribosomes Preferentially Translate Distinct Subpools of mRNAs Genome-wide. Molecular cell 67, 71-+, doi:10.1016/j.molcel.2017.05.021 (2017).

72 Gerst, J. E. Pimp My Ribosome: Ribosomal Protein Paralogs Specify Translational Control. Trends Genet 34, 832–845, doi:10.1016/j.tig.2018.08.004 (2018).

73 Pakos-Zebrucka, K. et al. The integrated stress response. EMBO reports 17, 1374–1395, doi:10.15252/embr.201642195 (2016).

74 Nandagopal, N. & Roux, P. P. Regulation of global and specific mRNA translation by the mTOR signaling pathway. Translation 3, e983402, doi:10.4161/21690731.2014.983402 (2015).

75 Merchante, C., Stepanova, A. N. & Alonso, J. M. Translation regulation in plants: an interesting past, an exciting present and a promising future (vol 90, pg 628, 2017). Plant Journal 98, 1157–1157, doi:10.1111/tpj.14359 (2019).

76 Kazama, T. et al. Curing cytoplasmic male sterility via TALEN-mediated mitochondrial genome editing. Nature plants 5, 722–730, doi:10.1038/s41477-019-0459-z (2019).

77 Kazama, T. et al. Curing cytoplasmic male sterility via TALEN-mediated mitochondrial genome editing. Nat Plants 5, 722–730, doi:10.1038/s41477-019-0459-z (2019).

78 Maliga, P. Engineering the plastid and mitochondrial genomes of flowering plants. Nat Plants, doi:10.1038/s41477-022-01227-6 (2022).

79 Arimura, S. I. et al. Targeted gene disruption of ATP synthases 6-1 and 6-2 in the mitochondrial genome of Arabidopsis thaliana by mitoTALENs. Plant J 104, 1459–1471, doi:10.1111/tpj.15041 (2020).

80 Nakazato, I. et al. Targeted base editing in the mitochondrial genome of Arabidopsis thaliana. Proc Natl Acad Sci U S A 119, e2121177119, doi:10.1073/pnas.2121177119 (2022).

81 Richter, U. et al. A Mitochondrial Ribosomal and RNA Decay Pathway Blocks Cell Proliferation. Current Biology 23, 535–541, doi:10.1016/j.cub.2013.02.019 (2013).

82 Gorgoulis, V. et al. Cellular Senescence: Defining a Path Forward. Cell 179, 813–827, doi:10.1016/j.cell.2019.10.005 (2019).

83 Mehlmer, N. et al. A toolset of aequorin expression vectors for in planta studies of subcellular calcium concentrations in Arabidopsis thaliana. Journal of experimental botany 63, 1751–1761, doi:10.1093/jxb/err406 (2012).

84 Knight, H., Trewavas, A. J. & Knight, M. R. Cold calcium signaling in Arabidopsis involves two cellular pools and a change in calcium signature after acclimation. The Plant cell 8, 489–503, doi:10.1105/tpc.8.3.489 (1996).

85 Mustroph, A., Juntawong, P. & Bailey-Serres, J. Isolation of Plant Polysomal mRNA by Differential Centrifugation and Ribosome Immunopurification Methods. Plant Systems Biology 553, 109–126, doi:10.1007/978-1-60327-563-7_6 (2009).

86 Missra, A. & von Arnim, A. G. Analysis of mRNA translation states in Arabidopsis over the diurnal cycle by polysome microarray. Methods in molecular biology 1158, 157–174, doi:10.1007/978-1-4939-0700-7_10 (2014).

87 Ma, J. et al. iProX: an integrated proteome resource. Nucleic Acids Res 47, D1211–D1217, doi:10.1093/nar/gky869 (2019).

88 Haug, K. et al. MetaboLights: a resource evolving in response to the needs of its scientific community. Nucleic Acids Res 48, D440–D444, doi:10.1093/nar/gkz1019 (2020).

