## Supplementary Figure 1-8 for "Perturbation of mitochondrial Ca^2+^ homeostasis activates cross-compartmental proteostatic response in Arabidopsis"

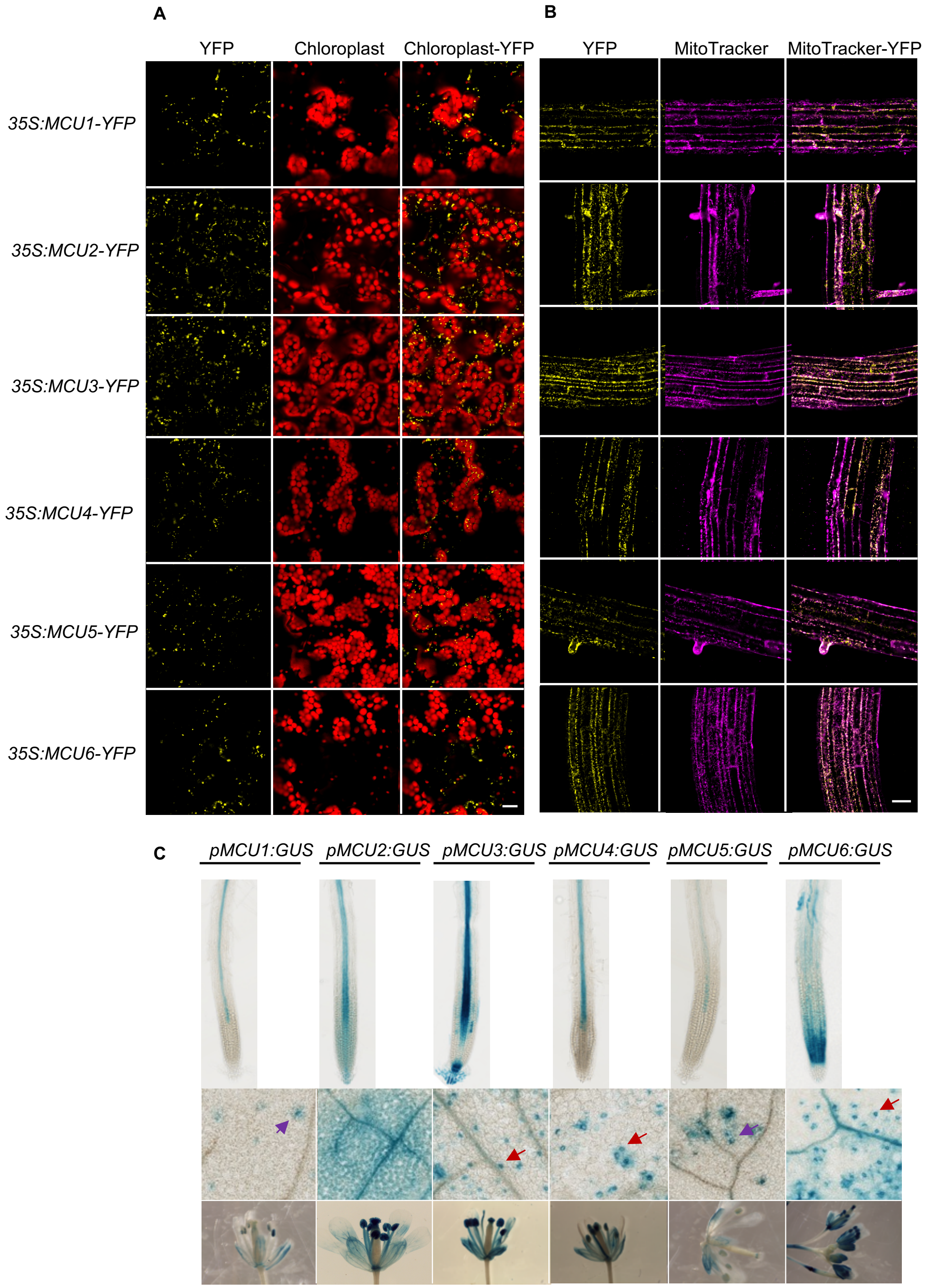


**Supplementary Fig. 1. Arabidopsis MCU proteins localize to mitochondria cross the tissues.** (A) YFP signals in the leaf epidermal and mesophyll cells of stable Arabidopsis transgenic plants expressing *35S:MCU-YFP* as indicated. Autofluorescence of chlorophyll is shown in red. (B) YFP signals in root cells of stable Arabidopsis transgenic plants expressing *35S:MCU-YFP* as indicated. The root was stained with MitoTracker™ Deep Red FM (Invitrogen M22426). YFP signals in root cells are merged with the red signals of MitoTracker. Scale bars, 20 μm. (C) GUS-staining highlights tissue expression of MCUs. Expression patterns of MCU genes are shown in roots (upper panel), leaves (middle panel) and flowers (lower panel). Red arrows indicate guard cells and purple arrows indicate trichrome.

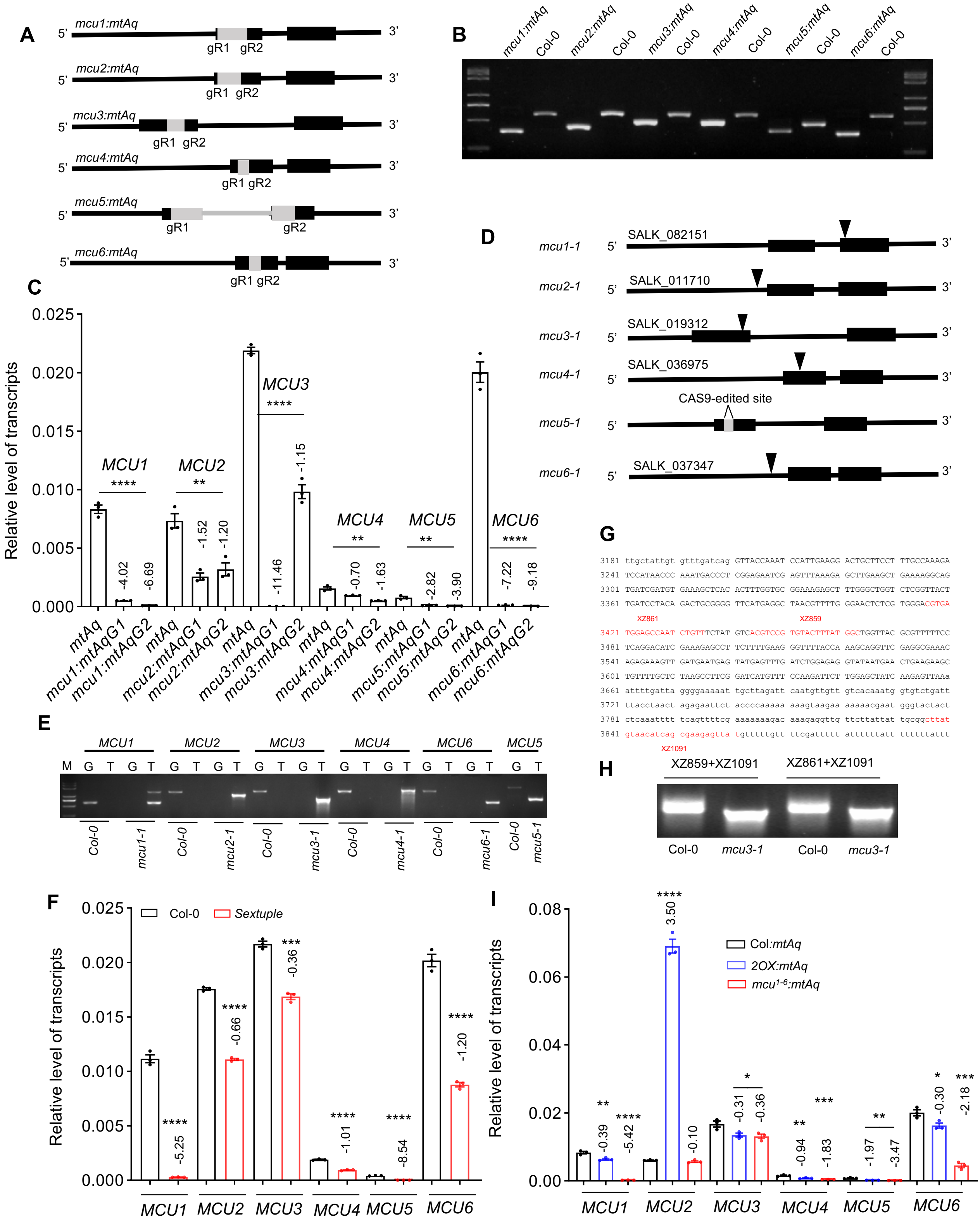
A
5' mcu1:mtAq 3'
gR1 gR2
5' mcu2:mtAq 3'
gR1 gR2
5' mcu3:mtAq 3'
gR1 gR2
5' mcu4:mtAq 3'
gR1 gR2
5' mcu5:mtAq 3'
gR1 gR2
5' mcu6:mtAq 3'
gR1 gR2
B
mcu1:mtAq Col-0 mcu2:mtAq Col-0 mcu3:mtAq Col-0 mcu4:mtAq Col-0 mcu5:mtAq Col-0 mcu6:mtAq Col-0
C
Relative level of transcripts
0.025
0.020
0.015
0.010
0.005
0.000
MCU1 **** -4.02 -6.69
MCU2 ** -1.52 -1.20
MCU3 **** -11.46 -1.15
MCU4 ** -0.70 -1.63
MCU5 ** -2.82 -3.90
MCU6 **** -7.22 -9.18
mtAq mcu1:mtAqG1 mcu1:mtAqG2 mtAq mcu2:mtAqG1 mcu2:mtAqG2 mtAq mcu3:mtAqG1 mcu3:mtAqG2 mtAq mcu4:mtAqG1 mcu4:mtAqG2 mtAq mcu5:mtAqG1 mcu5:mtAqG2 mtAq mcu6:mtAqG1 mcu6:mtAqG2
D
mcu1-1 5' SALK_082151 3'
mcu2-1 5' SALK_011710 3'
mcu3-1 5' SALK_019312 3'
mcu4-1 5' SALK_036975 3'
mcu5-1 5' CAS9-edited site 3'
mcu6-1 5' SALK_037347 3'
E
MCU1 MCU2 MCU3 MCU4 MCU6 MCU5
M G T G T G T G T G T G T G T G T G T G T G T
Col-0 mcu1-1 Col-0 mcu2-1 Col-0 mcu3-1 Col-0 mcu4-1 Col-0 mcu6-1 Col-0 mcu5-1
G
3181 ttgctattgt gtttgatcag GTTACCAAAT CCATTGAAGG ACTGCTTCCT TTGCCAAAGA
3241 TCCATAACCC AAATGACCCT CGGAGAATCG AGTTTAAAGA GCTTGAAGCT GAAAAGGCAG
3301 TGATCGATGT GAAAGCTCAC ACTTTGGTGC GGAAAGAGCT TTGGGCTGGT CTCGGTTACT
3361 TGATCCTACA GACTGCGGGG TTCATGAGGC TAACGTTTTG GGAACTCTCG TGGGACGTGA
XZ861 XZ859
3421 TGGAGCCAAT CTGTTTCTAT GTCACGTCCG TGTACTTTAT GGCTGGTTAC GCGTTTTTCC
3481 TCAGGACATC GAAAGAGCCT TCTTTTGAAG GGTTTTACCA AAGCAGGTTC GAGGCGAAAC
3541 AGAGAAAGTT GATGAATGAG TATGAGTTTG ATCTGGAGAG GTATAATGAA CTGAAGAAGC
3601 TGTTTTGCTC TAAGCCTTCG GATCATGTTT CCAAGATTCT TGGAGCTATC AAGAGTTAAa
3661 attttgatta ggggaaaaat tgcttagatt caatgttgtt gtcacaaatg gtgtctgatt
3721 ttacctaact agagaattct accccaaaaa aaagtaagaa aaaaacgaat gggtactact
3781 ctcaaatttt tcagttttcg aaaaaaagac aaagaggttg ttcttattat tgcggcttat
3841 gtaacatcag cgaagagtta tgtttttgtt ttcgattttt atttttattt tttttattt
XZ1091
H
XZ859+XZ1091 XZ861+XZ1091
Col-0 mcu3-1 Col-0 mcu3-1
F
Relative level of transcripts
0.025
0.020
0.015
0.010
0.005
0.000
Col-0 Sextuple
**** -5.25
**** -0.66
*** -0.36
**** -1.01
**** -8.54
**** -1.20
MCU1 MCU2 MCU3 MCU4 MCU5 MCU6
I
Relative level of transcripts
0.08
0.06
0.04
0.02
0.00
Col:mtAq
2OX:mtAq
mcu1-6:mtAq
** -0.39 **** -5.42
**** 3.50 -0.10
* -0.31 -0.36
** -0.94 *** -1.83
** -1.97 -3.47
* -0.30 *** -2.18
MCU1 MCU2 MCU3 MCU4 MCU5 MCU6

**Supplementary Fig. 2. Generation of *mcu* mutants and $Ca^{2+}$ reporter lines.** (A) Generation of CRISPR/Cas9-edited *mcu* mutants in a mitochondrion matrix-targeted $Ca^{2+}$ reporter line. Schematic diagrams show two sgRNA targeted (gR1 and gR2) sites within *MCU1*, *MCU2*, *MCU3*, *MCU4*, *MCU5*, and *MCU6* genes. (B) DNA gel image shows verification of a genomic DNA deletion (grey boxes indicated in A) in CRISPR/Cas9-edited single mutants by PCR. (C) Relative transcript levels of *MCU*s in CRISPR/Cas9-edited MCU single mutants. MCU transcripts were detected in two generations (G1 and G2) of CRISPR/Cas9-edited MCU single mutants using RT-qPCR. The numbers above the columns indicate log2 fold-changes in transcripts of mutants relative to the wildtype. (D) Identification of *mcu* T-DNA insertion mutants. Schematic diagrams show the positions of the T-DNA insertions within *MCU1*, *MCU2*, *MCU3*, *MCU4*, and *MCU6* and the CRISPR/Cas9-edited sites within *MCU5.* (E) The DNA gel image shows PCR confirmation of mutant lines. G stands for gene-specific primer pairs for detecting the presence of gene; and T stands for T-DNA specific primer pairs for detecting the presence of T-DNA. (F) Relative transcript levels of *MCU*s in sextuple mutant. MCU transcripts were detected in sextuple mutant using RT-qPCR. The numbers above the columns indicate log2 fold-changes in transcripts of sextuple mutants relative to the wild type. (G) Aberrant *MCU3* transcript in the *mcu3-1* mutant. Primers for RT-PCR are indicated in red on the *MCU3* sequence. (H) The sizes of *MCU3* transcripts in Col-0 and *mcu3-1* are shown in the gel image. (I). Relative transcript levels of *MCU*s in Col*:mtAq*, *2OX:mtAq* and $mcu^{1\text{-}6}$*:mtAq* lines. MCU transcripts were detected by RT-qPCR. The numbers above the columns indicate log2 fold-changes in transcripts of *2OX:mtAq* and $mcu^{1\text{-}6}$*:mtAq* relative to the wild type. Results are expressed as the mean ± SEM (n=3). Two-tailed student's *t*-test was performed to examine statistical significance for C, F and I, *$p < 0.05$, **$p < 0.01$, ***$p < 0.001$, ****$p<0.0001$. See also Supplementary Data11.

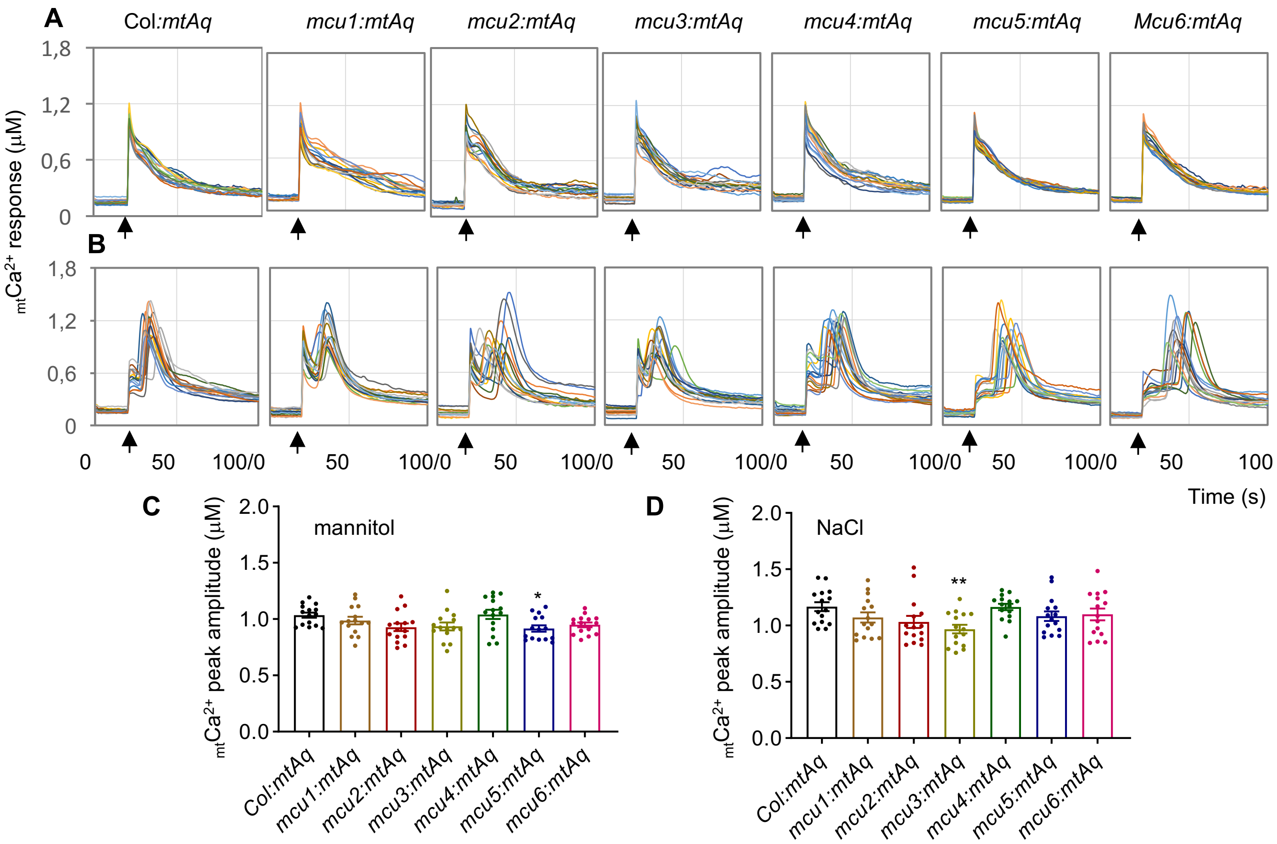


**Supplementary Fig. 3. Stress stimuli-induced mitochondrial $Ca^{2+}$ response is not significantly altered in *mcu* single mutants.** (A-B) The $_{mt}Ca^{2+}$ dynamic response. Aequorin-based $Ca^{2+}$ luminescence reading shows $_{mt}Ca^{2+}$ dynamic responses of wild-type and *mcu1*, *mcu2*, *mcu3*, *mcu4*, *mcu5*, and *mcu6* mutant seedlings subjected to 100 mM NaCl (A) or 400 mM mannitol (B). The luminescence was recorded at 1-s intervals, and the arrow indicates the starting point of the treatment. Each colored line represents the $_{mt}Ca^{2+}$ dynamic response of 1 of 15 individual seedlings. (C-D). The $_{mt}Ca^{2+}$ peak amplitudes. The $_{mt}Ca^{2+}$ luminescence peak value of 15 individual seedlings were determined by luminometer and converted to $_{mt}Ca^{2+}$ concentration as $_{mt}Ca^{2+}$ peak amplitudes (see methods). The $_{mt}Ca^{2+}$ peak amplitudes in response to mannitol (C) and NaCl (D) in *mcu1:mtAq*, *mcu2:mtAq*, *mcu3:mtAq*, *mcu4:mtAq*, *mcu5:mtAq*, and *mcu6:mtAq* single mutants are not significantly altered compared to the wild type. Results are expressed as the mean ± SEM (n=15). One-way ANOVA followed by Dunnett's multiple comparisons test was performed to determine significance, *$p < 0.05$, **$p < 0.01$. ***$p$<0.001, ****$p$<0.0001. See also Supplementary Data 11 for the information of statistical analysis.

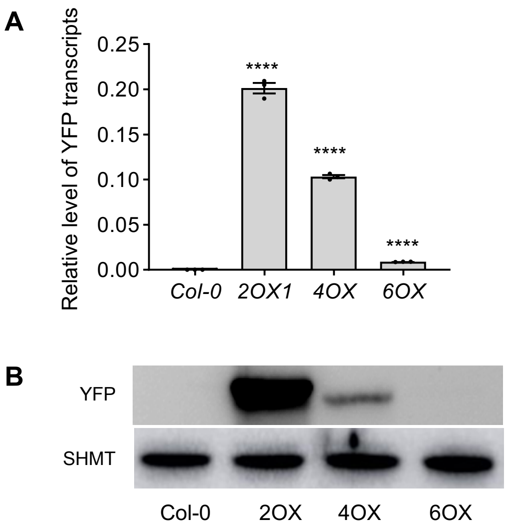


**Supplementary Fig. 4. MCU protein levels in *MCU* overexpression lines.** (A) Relative transcript levels of *MCU2*, *MCU4*, and *MCU6* in *2OX*, *4OX*, and *6OX* lines, determined from the relative transcript levels of *YFP* by RT-qPCR. Results are expressed as the mean ± SEM (n=3). Two-tailed student's *t*-test was performed to examine statistical significance, ****$p < 0.0001$. See also Supplementary Data 11. (B) Protein abundance of MCU2, MCU4, and MCU6 in *2OX*, *4OX*, and *6OX* lines, determined by immunoblotting with an anti-YFP antibody. SHMT was used as loading control. Ten-day-old seedlings grown vertically on MS plates were collected for the analysis.

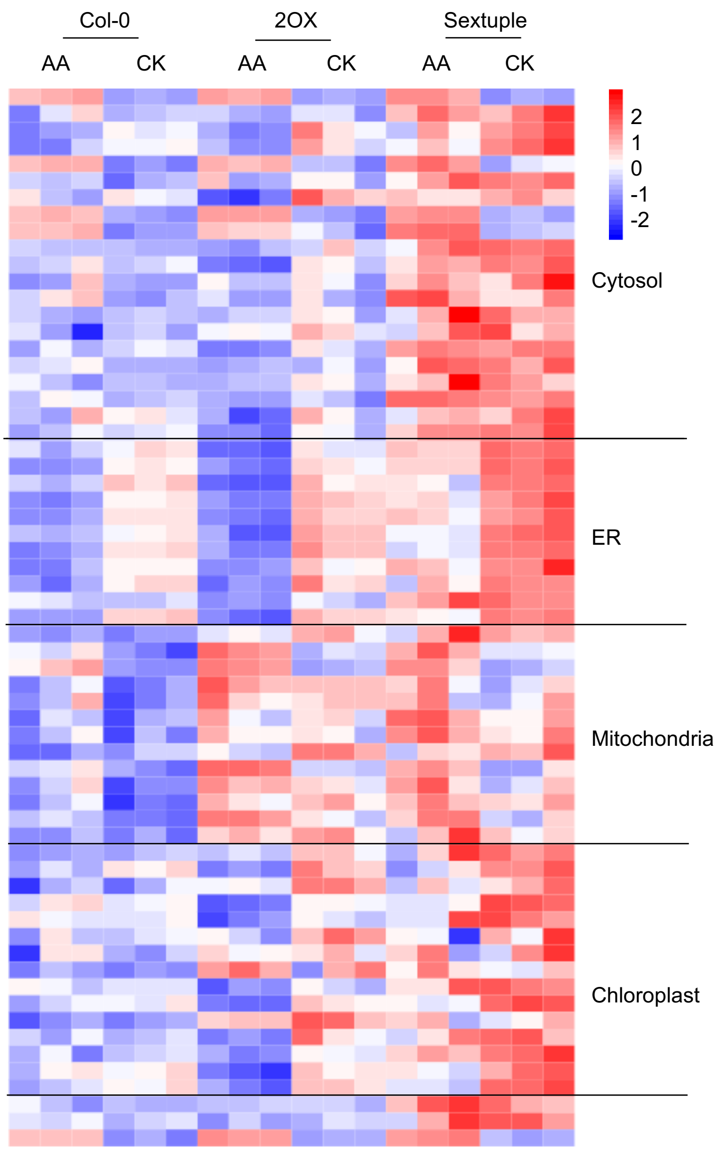


**Supplementary Fig. 5. Heatmap of differentially expressed chaperone genes.** Differentially expressed chaperone genes within different cellular compartments, as indicated by a heatmap in AA-treated samples compared to untreated controls. See also Supplementary Data 3 for the list of gene ID.

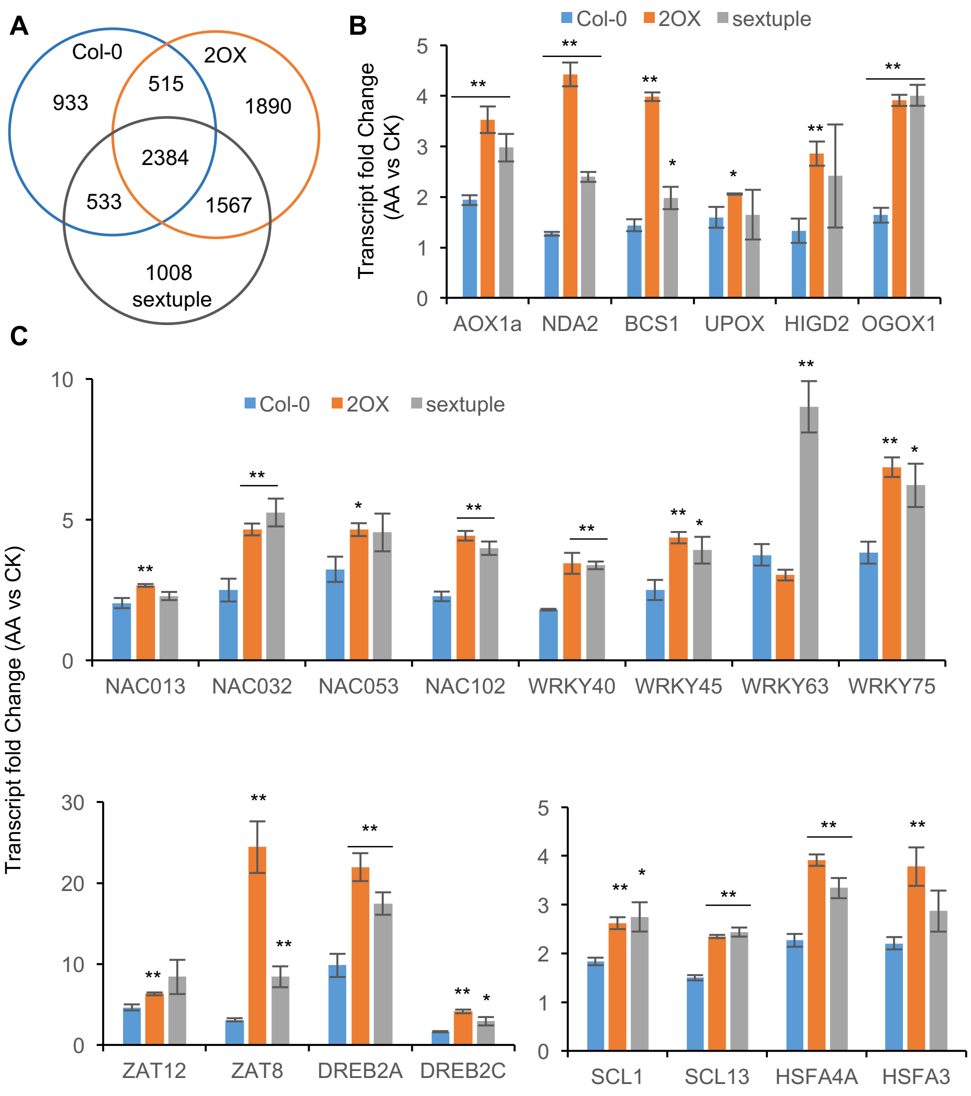


**Supplementary Fig. 6. Perturbation of $_{mt}Ca^{2+}$ homeostasis enhances AA-triggered mitochondrial retrograde response (MRR) and general stress response**. (A) Venn diagram of DEGs in the AA-treated wildtype, *2OX* and the sextuple mutant, compared to their controls, as determined by RNA-seq (Benjamini-Hochberg adjusted *p*-value < 0.01). See Supplementary Data 1 for a complete list of DEGs. (B-C) MRR marker (B) and TF (C) gene expression in wild-type, *2OX* and *mcu* sextuple mutant, expressed as fold-changes in AA-treated samples relative to their controls. Results are expressed as the mean ± SEM (n=3). Two-tailed student's *t*-test was performed to examine statistical significance, **p* < 0.05, ***p* < 0.01. See also Supplementary Data 11 for the information of statistical analysis.

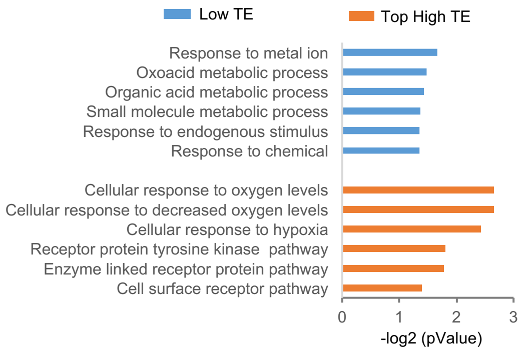


**Supplementary Fig. 7. GO enrichment of the genes with saTE in AA-treated wildtype. A** Top six biological processes with highest enrichment scores in low TE (blue) or high TE (yellow) transcripts for each GO category.

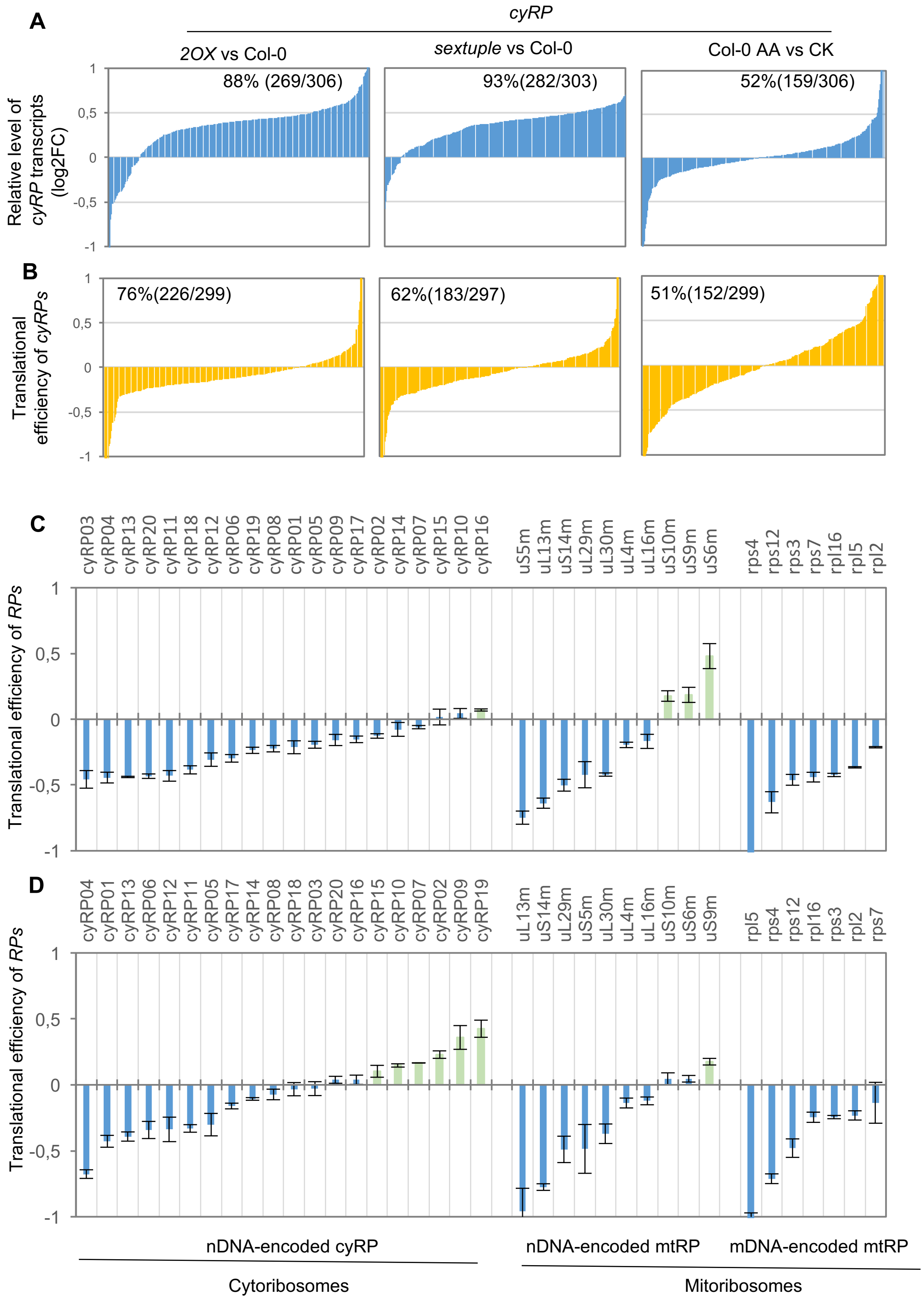


**Supplementary Fig. 8. Translational repression of *cyRPs*.** (A-B) Levels of individual *cyRP* transcripts and TEs are plotted, showing a large number of upregulated *cyRP* transcripts with low TE. The percentage of upregulated *cyRP* transcripts (A) and *cyPR* mRNAs with low TE (B) are indicated in each plot. (C-D) Verification of *cyRP* mRNA with low TE. TEs of cytoribosome and mitoribosome mRNAs were examined by RT-qPCR using RNA isolated from monosomal and polysomal fractions of *2OX* (C) and sextuple mutant (D).
